# Evolution of connectivity architecture in the *Drosophila* mushroom body

**DOI:** 10.1101/2023.02.10.528036

**Authors:** Kaitlyn Elizabeth Ellis, Sven Bervoets, Hayley Smihula, Ishani Ganguly, Eva Vigato, Thomas O. Auer, Richard Benton, Ashok Litwin-Kumar, Sophie Jeanne Cécile Caron

## Abstract

Brain evolution has primarily been studied at the macroscopic level by comparing the relative size of homologous brain centers between species. How neuronal circuits change at the cellular level over evolutionary time remains largely unanswered. Here, using a phylogenetically informed framework, we compare the olfactory circuits of three closely related *Drosophila* species that differ radically in their chemical ecology: the generalists *Drosophila melanogaster* and *Drosophila simulans* that feed on fermenting fruit, and *Drosophila sechellia* that specializes on ripe noni fruit. We examine a central part of the olfactory circuit that has not yet been investigated in these species — the connections between the projection neurons of the antennal lobe and the Kenyon cells of the mushroom body, an associative brain center — to identify species-specific connectivity patterns. We found that neurons encoding food odors — the DC3 neurons in *D. melanogaster* and *D. simulans* and the DL2d neurons in *D. sechellia* — connect more frequently with Kenyon cells, giving rise to species-specific biases in connectivity. These species-specific differences in connectivity reflect two distinct neuronal phenotypes: in the number of projection neurons or in the number of presynaptic boutons formed by individual projection neurons. Finally, behavioral analyses suggest that such increased connectivity enhances learning performance in an associative task. Our study shows how fine-grained aspects of connectivity architecture in an associative brain center can change during evolution to reflect the chemical ecology of a species.

## MAIN TEXT

Brain evolution has been primarily studied at the macroscopic level by comparing gross neuroanatomical features in homologous brain centers across distantly related species^1–3^. This pioneering work revealed that, over intermediate evolutionary timescales, the number of specialized brain centers does not change considerably, but the types and number of neurons forming these centers can vary greatly. Recent advances in comparative transcriptomics have provided new insights into the evolution and diversification of neurons, revealing how subtle variations in highly conserved regulatory gene networks can give rise to drastic changes in the rate at which neuronal progenitors proliferate or the types of neuron they give rise to^4^. To what degree such changes reflect evolutionary pressures remains unclear. Moreover, how such changes manifest themselves at the level of neuronal circuits is not yet understood, as it remains technically challenging to delineate neuronal circuits in non-traditional model systems and to compare them across species with different evolutionary trajectories^5^.

Flies in the genus *Drosophila* have evolved to exploit a remarkable diversity of ecological niches^6^. The phylogeny of most *Drosophila* species has been resolved, revealing that even closely related *Drosophila* species can inhabit drastically different environments^7^. For instance, *Drosophila sechellia*, a species endemic to the Seychelles islands, is a specialist for noni — a toxic fruit that produces a distinctive bouquet of pungent acids — whereas its closest relative, *Drosophila simulans*, is a generalist and a human commensal that can be found in most cosmopolitan areas^8–11^ (Figure 1a). A mere 0.25 million years of evolution separate these two species from their common ancestor, and only 3 million years separate their common ancestor from their well-studied relative, *Drosophila melanogaster*^7^*. D. melanogaster* is also a human commensal whose ecology largely overlaps with that of *D. simulans*^12^. These inverse relationships — phylogenetically, *D. sechellia* is closer to *D. simulans* while ecologically *D. melanogaster and D. simulans* are more alike — make this group of *Drosophila* highly suitable for investigating how neuronal circuits evolve over comparatively short timescales.

**Figure 1.**
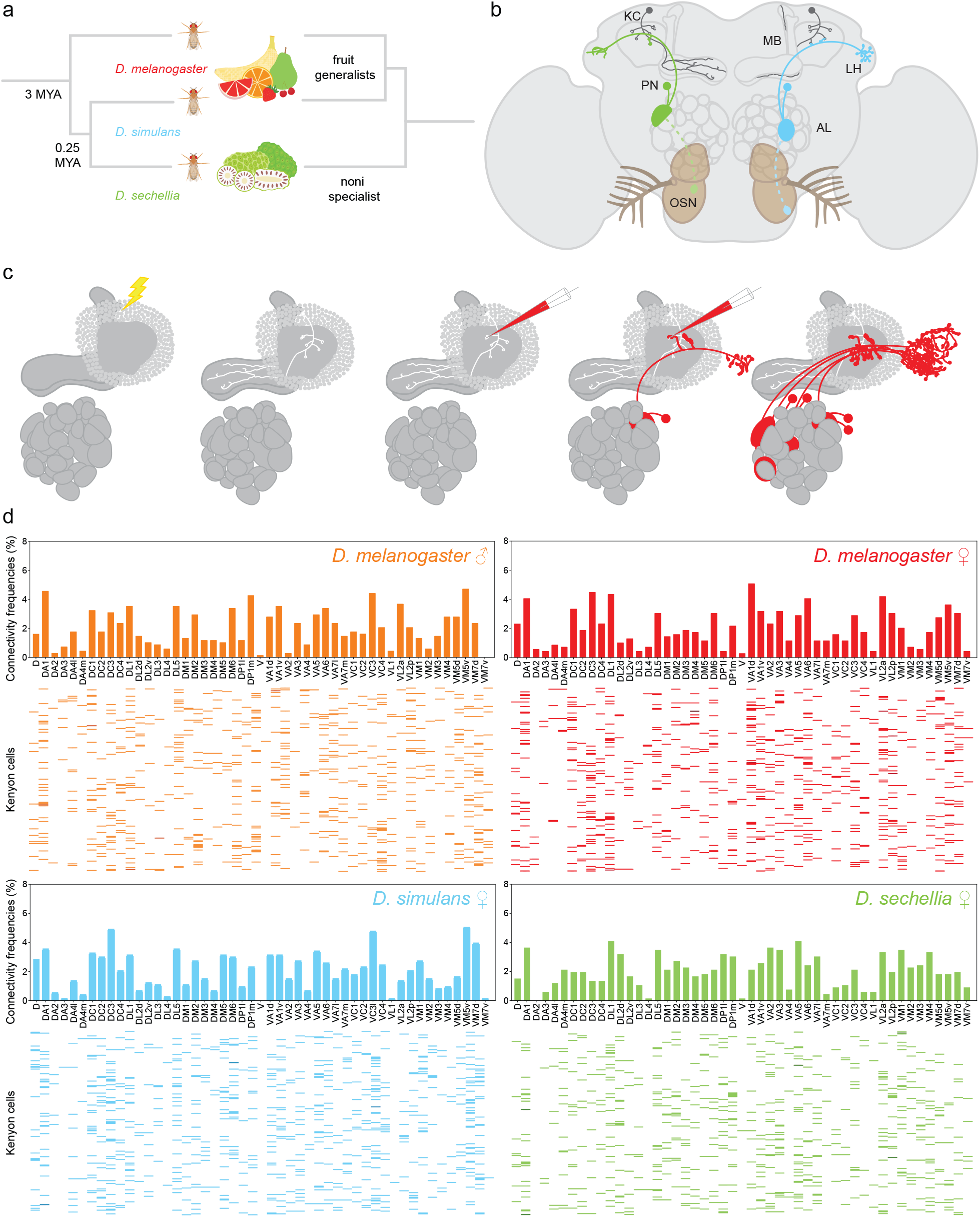
Mapping Kenyon cell inputs in *Drosophila* species living in different ecological niches. (a) Schematic depicting the phylogenetic relationships of *D. melanogaster* (red), *D. simulans* (blue), and *D. sechellia* (green) on the left and their ecological relationships on the right. (b) Schematic depicting the *Drosophila* olfactory circuit: olfactory sensory neurons that express the same receptor gene(s) (OSNs, green and blue neurons with dotted outline) converge onto the same glomerulus in the antennal lobe (AL); projection neurons (PNs, green and blue neurons with full outline) connect individual glomeruli to the mushroom body (MB) and the lateral horn (LH); Kenyon cells (dark grey) receive input from a small number of projection neurons. (c) Schematic depicting the technique used to map connections between projection neurons and Kenyon cells: a Kenyon cell is photo-labelled (white) and the projection neurons connected to each of its claw are dye-labeled (red) such that the antennal lobe glomeruli innervated by the labeled projection neurons can be identified; see Extended Figure 3 for a more detailed description of the technique. (d) Connections between glomeruli and Kenyon cells were mapped in *D. melanogaster*, *D. simulans* and *D. sechellia*, and all connections are reported in four connectivity matrices (*D. melanogaster* males: top left panel and orange (687 connections); *D. melanogaster* females: top right panel and red (704 connections); *D. simulans* females: bottom left panel and blue (717 connections); *D. sechellia* females: bottom right panel and green (692 connections)). In each matrix, a row corresponds to a Kenyon cell — there are 200 Kenyon cells per matrix — and each column corresponds to the different antennal lobe glomeruli; each colored bar indicates the input connections of a given Kenyon cell, and the intensity of the color denotes the number of connections found between a particular Kenyon cell and a given glomerulus (light: one connection; medium: two connections; dark: three connections). The bar graphs above the matrices represent the frequencies at which a particular glomerulus was connected to Kenyon cells as measured in a given matrix.

Olfactory driven behaviors used to locate food sources are most likely one of the most important ways whereby a species can adapt to its ecological niche. In *D. melanogaster,* most olfactory sensory neurons express only one receptor gene, and the axons of neurons expressing the same receptor gene(s) converge on a specific glomerulus in the antennal lobe, forming a stereotypical map^13,14^ (Figure 1b). The genomes of *D. melanogaster*, *D. simulans* and *D. sechellia* harbor a comparable number of functional olfactory receptor genes, and homologous glomeruli can be identified in the antennal lobe in these species^15,16^. In *D. sechellia*, however, a few olfactory sensory neurons show species-specific responses to noni odors: Or22a-expressing neurons are preferentially activated by methyl esters whereas Ir75b-expressing neurons are most sensitive to hexanoic acid^16–20^. In *D. melanogaster,* Or22a-expressing neurons are broadly tuned to different ethyl esters whereas Ir75b-expressing neurons are broadly tuned to shorter-chain acids^16–20^. These species-specific tuning properties are due to specific amino acid differences in the presumed ligand-binding domain of Or22a and Ir75b receptors^16,20^. In addition, there are two-to three-fold more Or22a-and Ir75b-expressing neurons in *D. sechellia*, and the glomeruli innervated by these neurons, DM2 and DL2d, respectively, are larger^16–18^. Other glomeruli — such as VM5d, which is innervated by Or85c/b-expressing neurons — are also larger in *D. sechellia* compared to *D. melanogaster*^16,18,19^

Whether and how the changes that occurred at the levels of receptor proteins and sensory neurons are reflected downstream, at the level of higher processing centers, is largely unknown. Projection neurons innervating individual antennal lobe glomeruli relay olfactory information to the mushroom body, an associative brain center, and the lateral horn, a center that mediates innate responses to odors^21^ (Figure 1b). The projection neurons of the antennal lobe and the Kenyon cells of the mushroom body have been studied in detail in *D. melanogaster*. Each glomerulus type is innervated by a distinct, but largely stereotyped number of projection neurons, from one up to eight^22^. The mushroom body consists of about 2,000 neurons, called ‘Kenyon cells’, that can be divided into three major types (α/β, α’/β’ and γ Kenyon cells); each Kenyon cell receives input from a small number of projection neurons, on average seven^23,24^. The connectivity architecture between projection neurons and Kenyon cells has been resolved, revealing two important principles: first, these connections are unstructured, in that individual Kenyon cells integrate inputs from a random set of projection neurons; second, some types of projection neuron connect more frequently to Kenyon cells than others, leading to a biased representation of glomeruli in the mushroom body^24–27^. These two connectivity patterns — randomization of input and biased connectivity — are genetically hardwired, suggesting that they might be shaped by evolutionary pressures^28^.

## RESULTS

### Randomization of input is conserved across *Drosophila* species

To compare features of the connectivity architecture between projection neurons and Kenyon cells in *D. melanogaster*, *D. simulans* and *D. sechellia*, we first resolved gross anatomical features of the antennal lobes of these species. Using confocal images of brains immuno-stained with a neuropil marker, we reconstructed entire antennal lobes by manually tracing the borders of individual glomeruli on single planes and projecting their volumes in three-dimensional space (Extended Data Figure 1). The resulting projections preserved the shape, volume and location of all the glomeruli forming an antennal lobe. Each glomerulus was annotated using the well-characterized *D. melanogaster* antennal lobe map, which was generated using similar methods, as a reference^13,14,29^. Glomerular volumes were compared across species (Extended Data Table 1). We found that the antennal lobe map is largely conserved in the three species and contains a total of 51 glomeruli that can be recognized based on their shape and location. The volumes of most glomeruli are comparable across species with a few exceptions, including the noni-responsive DL2d, DM2 and VM5d glomeruli, which are larger in *D. sechellia*, as previously reported (DL2d: *D. melanogaster*: 1565 ± 334 μm^3^ (*n* = 3); *D. simulans*: 2007 ± 185 μm^3^ (*n* = 3); *D. sechellia*: 3066 ± 296 μm^3^ (*n* = 3); DM2: *D. melanogaster*: 3139 ± 227 μm^3^ (*n* = 3); *D. simulans*: 3169 ± 227 μm^3^ (*n* = 3); *D. sechellia*: 4594 ± 127 μm^3^ (*n* = 3); VM5d: *D. melanogaster*: 905 ± 115 μm^3^ (*n* = 3); *D. simulans*: 1773 ± 127 μm^3^ (*n* = 3); *D. sechellia*: 3793 ± 153 μm^3^ (*n* = 3))^16,18,20^. Despite these volume differences, the antennal lobes of the three species investigated are macroscopically nearly identical, which is in line with the notion that gross anatomy is conserved over the fairly short evolutionary distances separating the three species.

**Table 1.** Morphological features of photo-labeled projection neurons across species. Morphological features of projection neurons — namely the number of neurons associated with a given glomerulus and the volume of the presynaptic sites or boutons these neurons form in the mushroom body — were measured and compared across species (*n* = 5 for each type of projection neuron, standard deviation is shown). Projection neurons showing significant shifts in connectivity frequencies (DC3, DL2d, VM5v, DP1l, VA1d, VC3, and VL2a projection neurons) were analyzed as well as some projection neurons that did not show significant shifts in connectivity frequencies (DA2, DM2, and DP1m projection neurons). Projection neurons were typed based on whether their cell bodies are located in the anterior-dorsal (ad), lateral (l) or ventral cluster (v).

| Projection neurons | Neuron number | | | Bouton cluster volume $\mu\text{m}^3$ | | |
| --- | --- | --- | --- | --- | --- | --- |
|  | *D. melanogaster* | *D. simulans* | *D. sechellia* | *D. melanogaster* | *D. simulans* | *D. sechellia* |
| DA2 | 4 ± 0.80 IPNs | 4 ± 1.02 IPNs | 4 ± 0.40 IPNs | 112.79 ± 11.30 | 110.95 ± 37.06 | 179.18 ± 31.76 |
| DC3 | 3 ± 0.29 adPNs | 3 adPNs, 3 vPNs | 2 ± 0.29 adPNs | 370.15 ± 195.68 | 364.06 ± 146.60 | 233.02 ± 83.80 |
| DL2d | 5 adPNs, 1 vPN | 5 adPNs, 1 vPN | 5 adPNs, 3 vPNs | 112.18 ± 12.41 | 104.44 ± 25.93 | 348.29 ± 186.63 |
| DM2 | 2 IPNs | 2 IPNs | 2 IPNs | 367.38 ± 54.87 | 309.52 ± 89.42 | 365.40 ± 52.39 |
| DP1l | 1 IPN, 1 vPN | 1 IPN, 2 vPNs | 1 IPN, 2 vPNs | 244.39 ± 69.00 | 232.41 ± 100.73 | 388.91 ± 75.88 |
| DP1m | 1 adPN | 1 adPN | 1 adPN | 221.59 ± 72.28 | 172.85 ± 113.80 | 260.95 ± 105.19 |
| VA1d | 3 adPNs, 1 vPN | 3 adPNs, 1 vPN | 3 ± 0.40 adPNs, 1 vPN | 205.89 ± 26.65 | 195.78 ± 109.79 | 155.87 ± 50.25 |
| VC3 | 3 adPNs | 3 adPNs | 3 adPNs | 200.70 ± 97.03 | 158.48 ± 90.40 | 143.59 ± 46.17 |
| VL2a | 1 adPN, 3 vPNs | 1 adPN, 3 vPNs | 1 adPN, 3 vPNs | 174.36 ± 64.22 | 156.05 ± 79.93 | 208.92 ± 60.55 |
| VM5v | 3 ± 0.49 adPNs | 3 ± 0.40 adPNs | 2 ± 0.33 adPNs | 256.78 ± 104.09 | 268.43 ± 132.67 | 140.14 ± 77.76 |

We next set out to compare the global connectivity architecture of the mushroom body across the three species by adapting the technique we previously developed to map projection neuron–Kenyon cell connections in *D. melanogaster*^25^. For each species, individual Kenyon cells were photo-labeled in flies carrying the broad neuronal driver *nSynaptobrevin-GAL4* and a *UAS-photoactivatable-GFP* effector transgene. In *D. melanogaster*, there are three major types of Kenyon cell — α/β, α’/β’ and γ Kenyon cells — and Kenyon cells from each type form a variable number of claw-shaped dendritic terminals. α/β, α’/β’ and γ Kenyon cells were found in all three species, appear morphologically indistinguishable from one another and form on average a comparable number of claw-shaped dendritic terminals^23^ (Extended Data Figure 2). To identify the projection neurons connected to a photo-labeled Kenyon cell, a red-dye was electroporated sequentially in most of the claw-shaped dendritic terminals formed by that cell (Figure 1c, Extended Data Figure 3). Using this technique, the inputs of hundreds of Kenyon cells were identified in terms of the glomeruli from which they originate. We reported these results in a connectivity matrix that summarizes the glomerular inputs to 200 Kenyon cells. Statistical analyses of the resulting matrix can be used to reveal structured patterns of connectivity, such as whether groups of glomeruli are preferentially connected to the same Kenyon cells or whether projection neuron–Kenyon cell connections are random and biased^25,28^. We generated a total of four such connectivity matrices (Figure 1d): two using *D. melanogaster* males and females (a total of 687 and 704 connections, respectively), one using *D. simulans* females (717 connections) and one using *D. sechellia* females (692 connections).

**Figure 2.**
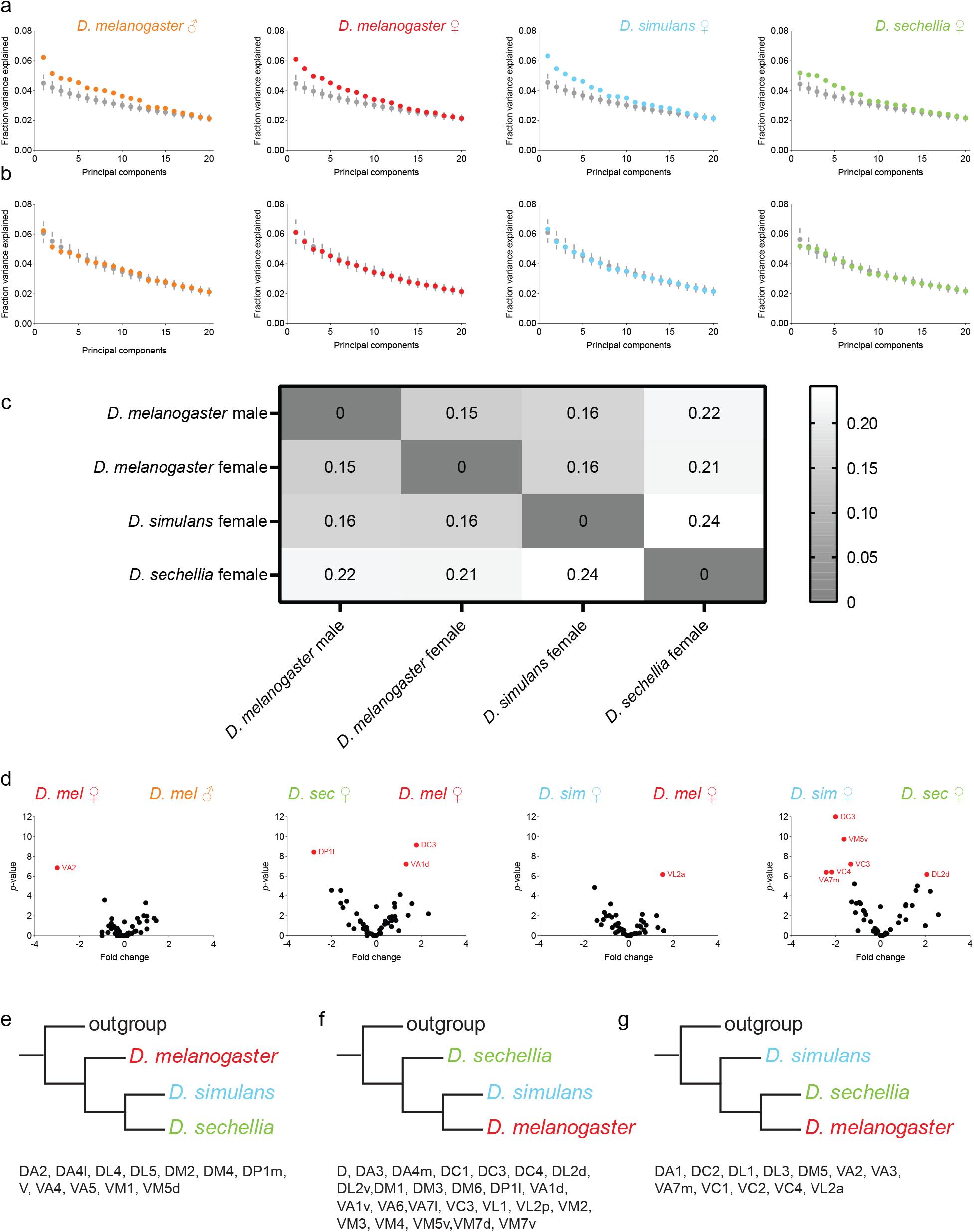
Shifts in connectivity biases across *Drosophila* species. (a-b) Principal components were extracted using each connectivity matrix as well as their uniform shuffle versions (a) or biased shuffle versions (b); the fraction of the variance explained by each component was measured (*D. melanogaster* males: far left panels and orange; *D. melanogaster* females: middle left panels and red; *D. simulans* females: middle right panels and blue; *D. sechellia* females: far right panels and green; colored circles: experimental matrices, grey circles: fixed-shuffle matrices); error bars represent 95% confidence interval. (c) The Jensen-Shannon distances were measured by comparing the distributions in connectivity frequencies observed experimentally and reported as a heat map; the color bar denotes the length of the distances measured. See Extended Data Figure 5 for the complete set of Jensen-Shannon distances including those measured when comparing the experimental matrices to their shuffle versions or when comparing the experimental matrices to an experimental matrix obtained in a previous study^28^. (d) The *p*-value measured for each type of projection neuron was plotted against the log_2_ fold change measured when comparing the connectivity frequencies measured for that type of projection neuron in the two matrices indicated on the plot. The statistical significance, or ‘*p*-value’, was measured for each type of projection neuron using the Fisher’s exact test; to control for false positives, *p*-values were adjusted with a false discovery rate of 0.10 using a Benjamini-Hochberg procedure. Within these plots, a fold change with a value of 0 indicates that there is no shift in frequencies between the two matrices, whereas a fold change that is smaller or greater than 0 indicates that a given type of projection neuron connects more frequently in one matrix than the other. Data points with *p*-values smaller than 0.01 are identified with a label (red circles); all other data points have *p*-values greater than 0.01 (black circles). (e-g) Similarity trees were generated using the Tree analysis Using New Technology (TNT) software. Trait used was the connectivity frequency measured for a given glomerulus in a given species (e-g) Trees generated for individual glomeruli were found to either follow the known phylogenetic relationships (e), the ecological relationships (f) or neither (g). See Extended Data Figure 6 for individual trees and more detailed explanations of the analyses.

We performed an unbiased search for potential structural features in these connectivity matrices using principal component analysis. The variance associated with individual principal component projections provides a sensitive measure of structure as we have previously shown^25,28^. We extracted correlations within a given experimental connectivity matrix and compared them to correlations extracted from matrices in which connections were randomly shuffled. In these shuffled matrices, the total number of connections between projection neurons and Kenyon cells is the same as the number of connections reported in each experimental matrix, but the connections were randomly assigned (’uniform shuffle matrices’). We also generated a second set of shuffle matrices in which the connections were scrambled but the frequencies at which projection neurons from each glomerulus connect to Kenyon cells were fixed to reflect the frequencies measured experimentally (’biased shuffle matrices’). We found that the observed spectrum of variances is not significantly different to that of the biased shuffle matrices, but does significantly deviate from that of the uniform shuffle matrices (Figure 2a,b). These results suggest that there are no detectable structural features in the experimental matrices other than the structure generated by the biases in connectivity frequencies.

### Biases in connectivity correlate with the chemical ecology of a species

If the connections between projection neurons and Kenyon cells were completely random, we would expect Kenyon cells to integrate input uniformly across glomeruli, and each glomerulus to have a connectivity frequency of about 2%. Yet, we found that the connections between projection neurons and Kenyon cells are biased: some glomeruli are overrepresented and have a connectivity frequency significantly higher than 2%, whereas some glomeruli are underrepresented and have a connectivity frequency significantly lower than 2% (Extended Data Figure 4). The result is a non-uniform distribution of connectivity frequencies. We found that there are between nine and 12 overrepresented glomeruli (glomeruli with connectivity frequencies higher than 2%, *p*-value < 0.05) and between 10 and 13 underrepresented glomeruli (glomeruli with connectivity frequencies lower than 2%, *p*-value < 0.05) within each connectivity matrix. This result shows that the non-uniform distribution of connectivity frequencies is a feature present across species.

To compare the overall extent of bias in the distributions of connectivity frequencies derived from the connectivity matrices, we inferred their Jensen-Shannon distances. This statistical method measures the similarity of two probability distributions, with a distance of zero indicating identical distributions. To gauge the extent to which the Jensen-Shannon distance indicates overall similarity of the observed distributions in connectivity frequencies, we compared the distributions of connectivity frequencies measured in the experimental matrices to those measured using the corresponding uniform shuffle matrices and obtained distances ranging from 0.23 to 0.27; when we compared the distributions of connectivity frequencies measured in the experimental matrices to those measured using the biased shuffle matrices, we obtained distances ranging from 0.08 to 0.09 (Extended Data Figure 5). When we compared the distributions of connectivity frequencies measured using two *D. melanogaster* female matrices — one matrix was generated in this study and the other was generated in a previous study^28^ — we obtained a relatively short distance of 0.17 (Figure 2c). Likewise, we obtained a relatively short distance of 0.15 when we compared the distributions measured in *D. melanogaster* females and males. This result suggests that the biases in connectivity are largely similar in both sexes. The distances measured when comparing *D. melanogaster* and *D. simulans* range from 0.16 to 0.17, indicating that the overall extent of bias in connectivity is similar in these species. By contrast, the distances between *D. melanogaster* and *D. sechellia* range from 0.20 to 0.22, whereas the distance between *D. simulans* and *D. sechellia* is 0.24, showing that the overall extent of biases in connectivity is higher in *D. sechellia* than in its sibling species. As *D. sechellia* and *D. simulans* are phylogenetically more closely related to each other than to *D. melanogaster*, the observed pattern suggests that the overall similarity in biases is not a function of evolutionary relatedness.

We next investigated whether the larger Jensen-Shannon distances measured for *D. sechellia* result from small differences in connectivity frequencies distributed across glomeruli or whether they result from large differences restricted to a few glomeruli. We performed pairwise comparisons and measured the ratio of connectivity frequencies obtained for a given glomerulus in two different matrices (connectivity frequency measured in matrix 1 divided by the connectivity frequency measured in matrix 2) (Figure 2d). We found that most glomeruli are connected to Kenyon cells at similar frequencies in the six possible pairwise comparisons but that nine glomeruli are connected at significantly different frequencies in at least one pairwise comparison (*p*-value < 0.01; Benjamini-Hochberg procedure with false discovery rate of 0.10, see Extended Data Table 2 for exact *p*-values). This result indicates that significant changes in only a fraction of the glomeruli account for the observed shift in biases between the three species.

In only one of the nine cases, the change concerned *D. simulans*, where the VL2a glomerulus was found to be connected at lower frequency than in *D. melanogaster* (VL2a connectivity frequencies: *D. simulans*: 1.36%, *D. melanogaster*: 4.21%, *p*-value = 0.002068546, *D. sechellia*: 3.34%, *p*-value = 0.044418781). In all other cases, the significant changes occurred in *D. sechellia*. Two glomeruli — DL2d and DP1l, which receive input from the acid-sensing Ir75b-and Ir75a-expressing neurons, respectively — were found to be connected at higher frequencies in *D. sechellia* than in one of the other two species (DL2d connectivity frequencies: *D. sechellia*: 3.19%, *D. simulans*: 0.68%, *p*-value = 0.002030026, *D. melanogaster*: 1.02%, *p*-value = 0.010694533; DP1l: *D. sechellia*: 3.19%, *D. melanogaster*: 0.44%, *p*-value = 0.000213936, *D. simulans*: 0.95%, *p*-value = 0.010694533). Six glomeruli — namely the DC3, VA1d, VA7m, VC3, VC4 and VM5v glomeruli — were connected at lower frequencies in *D. sechellia* than in one or both species. Four of these glomeruli receive input from olfactory sensory neurons tuned to various fruit volatiles: the Or83c-expressing neurons associated with the DC3 glomerulus are narrowly tuned to farnesol, a yeast odor, whereas the neurons associated with the VC3 (Or35a-expressing neurons), VC4 (Or67c-expressing neurons) and VM5v (Or98a-expressing neurons) glomeruli are broadly tuned to alcohols and esters produced by fermenting fruits^13,14,30–32^ (connectivity frequencies: DC3: *D. sechellia*: 1.37%, *D. melanogaster*: 4.50%, *p*-value = 0.000105671, *D. simulans*: 4.91%, *p*-value = 6.18E-0.6; VC3: *D. sechellia*: 2.13%, *D. simulans*: 5.00%, *p*-value = 0.000724388, *D. melanogaster*: 2.90%, *p*-value = 0.0.282487997; VC4: *D. sechellia*: 0.61%, *D. simulans*: 2.46%*, p*-value = 0.044418781, *D. melanogaster*: 1.74%, *p*-value = 0.0.040115307; VM5v: *D. sechellia*: 1.82%, *D. simulans*: 5.05%, *p*-value = 5.83E-0.5, *D. melanogaster*: 3.36%, *p*-value = 0.016285553). The olfactory sensory neurons associated with the VA1d glomerulus (Or88a-expressing neurons) are tuned to methyl palmitate, a pheromone made by *D. melanogaster* females but not by *D. simulans* or *D. sechellia* females^33^ (VA1d connectivity frequencies: *D. sechellia*: 2.13%, *D. melanogaster*: 5.08%, *p*-value = 0.000724388, *D. simulans*: 3.14%, *p*-value = 0.0.087663066). The receptor identity and cognate odors of the olfactory sensory neurons that connect to the VA7m glomerulus remain to be identified.

Overall, these results show that species-specific differences in connectivity frequencies are restricted to a fraction of olfactory channels, which relay information about pheromones and food odors to the mushroom body, suggesting that the representation of specific channels changed as these species diverged. Most notably, in all but one case, the connectivity frequency measured for a given glomerulus changes in the *D. sechellia* lineage, being more similar in *D. melanogaster* and *D. simulans* than when comparing any of these species to *D. sechellia*. As *D. sechellia* is more closely related to *D. simulans* than to *D. melanogaster*, this pattern appears not to be a function of phylogenetic relatedness, but instead of ecological relatedness. To better quantify this notion, we built a character matrix based on the connectivity frequencies measured for each glomerulus. We derived a similarity tree for each glomerulus such that we could determine whether the resulting tree reflects phylogenetic relationships or whether it reflects ecological relationships. We found that 26 trees, as well as the tree generated using the entire character matrix, follow the ecological relationships and that only 12 trees follow the phylogenetics relationships (Figure 2e-g, Extended Data Figure 6). To contrast this finding, we also derived a similarity tree using the coding sequence of the olfactory receptor(s) associated with a given glomerulus. Not surprisingly, we found that all these trees follow the phylogenetic relationships (Extended Data Figure 7). These observations show that changes in glomerular representation in the mushroom body are highly correlated with the ecology of a species, not its phylogeny, and, therefore, could have evolved in *D. sechellia* as this species diverged to exploit noni fruit.

### Two morphological features of projection neurons underlie shifts in biases

From our analyses of the above data sets, we observed two types of change in the mushroom body connectivity architecture of *D. sechellia* when compared to that of *D. melanogaster* and *D. simulans*: the representation of a few glomeruli either increases, for instance the representation of the DL2d and DP1l glomeruli, or decreases, for instance the representation of the DC3, VA1d, VA7m, VC3, VC4 and VM5v glomeruli. In *D. melanogaster*, it is known that biases in connectivity are a function of the overall number of presynaptic boutons formed in the mushroom body by the projection neurons associated with a given glomerulus^24,25,27^. Thus, changes in glomerular representation across species could result from changes in the number of projection neurons associated with a given glomerulus, changes in the number of presynaptic boutons individual projection neurons form or both. To determine whether any of these cases prevail, we photo-labeled (Figure 3a) and dye-labeled (Figure 4a) the projection neurons innervating a given glomerulus to quantify the number of neurons and measure the volume of the presynaptic boutons the labeled neurons occupy in the mushroom body.

**Figure 3.**
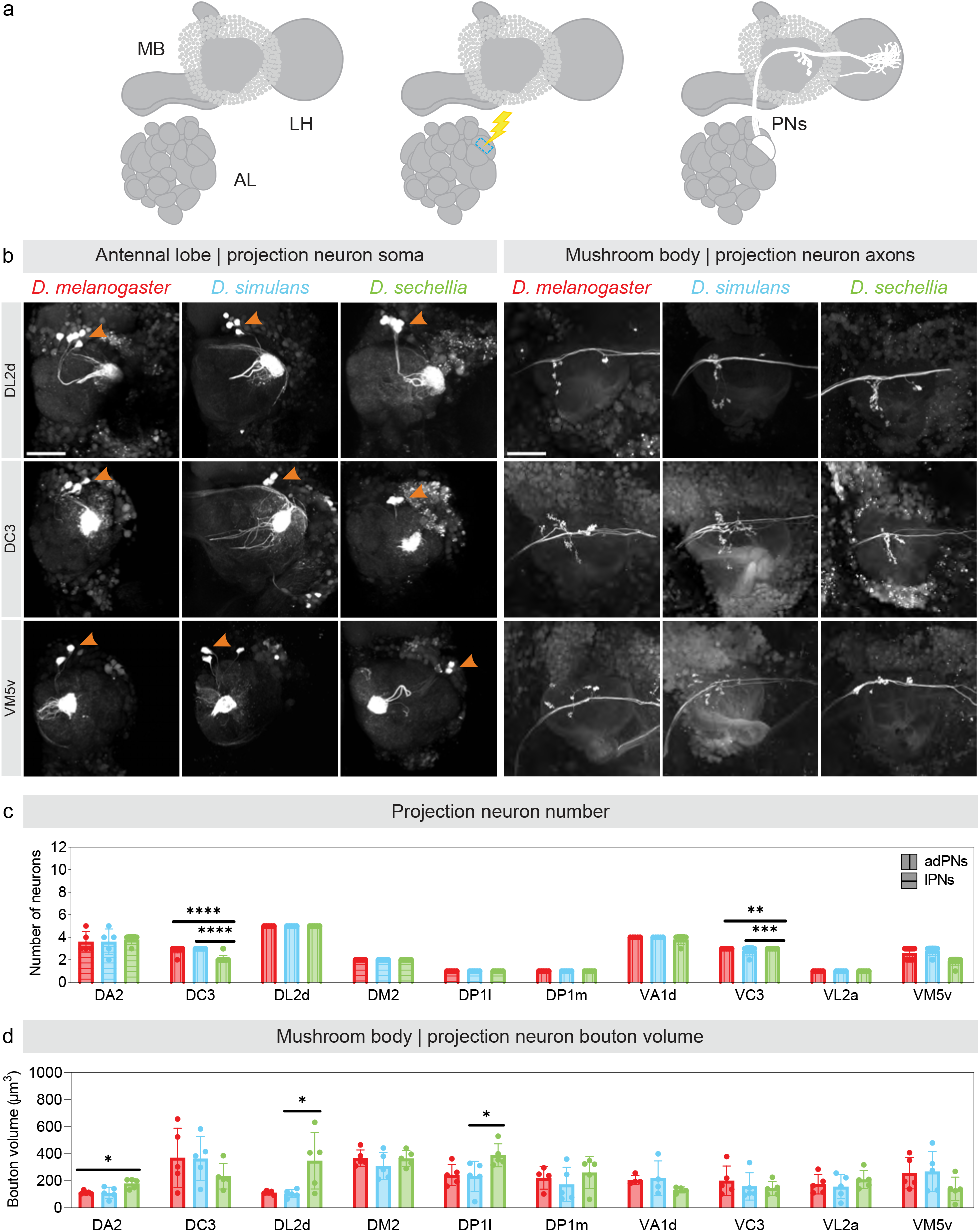
Morphological features of the projection neurons innervating a given glomerulus in different *Drosophila* species. (a) Schematic depicting the technique used to photo-label the projection neurons innervating a given glomerulus: a glomerulus is used as a landmark for photo-labeling (blue dashed outline), and the projection neurons connected to the targeted glomerulus are photo-labelled after successive rounds of photo-labeling. (b) The projection neurons innervating the DL2d (upper panels), DC3 (middle panels) and VM5v (lower panels) glomeruli were photo-labeled in *D. melanogaster* (left column of each panel), *D. simulans* (middle column of each panel) and *D. sechellia* (right column of each panel); the cell bodies of these neurons (left panels) and the axonal termini that these neurons extend in the mushroom body (right panels) were imaged. Scale bar is 50 µm. (c-d) The number of photo-labeled neurons (c) and the volume of the presynaptic boutons these neurons form in the mushroom body (d) were quantified and compared across species (red: *D. melanogaster*; blue: *D. simulans*; green: *D. sechellia*). The statistical significance, or ‘*p*-value’, was measured using the Mann-Whitney U test (*: *p*-value < 0.5, **: *p*-value < 0.01, ***: *p*-value < 0.001). See Table 1 for quantifications.

**Figure 4.**
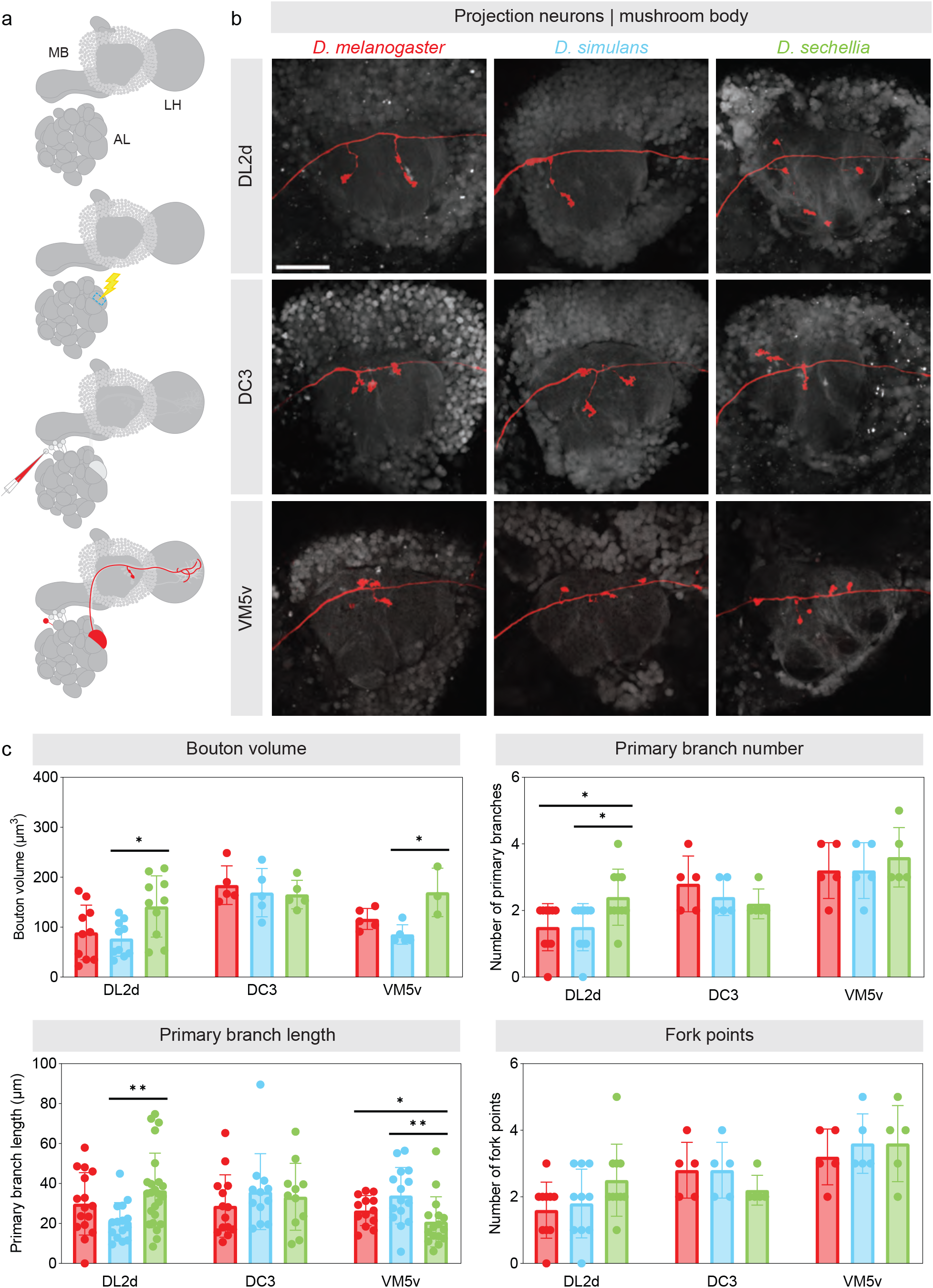
Morphological features of individual projection neurons in different *Drosophila* species. (a) Schematic depicting the technique used to dye-label a projection neuron innervating a given glomerulus: a glomerulus is used as a landmark for a first round of photo-labeling (blue dashed outline) during which the projection neurons connected to the targeted glomerulus are lightly photo-labelled; dye is electroporated in one of the photo-labeled projection neurons such that a single projection neuron is dye-labeled. (b) A projection neuron innervating the DL2d (upper row), DC3 (middle row) and VM5v (lower row) glomeruli were dye-labeled in *D. melanogaster* (left column), *D. simulans* (middle column) and *D. sechellia* (right column); the axonal termini these neurons extend in the mushroom body were imaged. Scale bar is 50 µm. (c-d) Various morphological features displayed by projection neurons in the mushroom body were quantified and compared across species (red: *D. melanogaster*; blue: *D. simulans*; green: *D. sechellia*). The statistical significance, or ‘*p*-value’, was measured using the Mann-Whitney U test (*: *p*-value < 0.5, **: *p*-value < 0.01, ***: *p*-value < 0.001). See Extended Data Table 3 for quantifications.

The DL2d and DP1l glomeruli are more frequently connected to Kenyon cells in *D. sechellia* (Figure 2d, Extended Data Table 2). We identified eight DL2d projection neurons in *D. sechellia* (five neurons located in the anterior-dorsal cluster, or adDL2d neurons, and three neurons located in the ventral cluster, or vDL2d neurons) but only six DL2d projection neurons in *D. melanogaster* and *D. simulans* (five adDL2d neurons and one vDL2d neuron in both species) (Table 1, Figure 3b-c). The number of DL2d projection neurons identified in *D. melanogaster* is consistent with the number of projection neurons reported in the available connectomes of the *D. melanogaster* brain^22^ (Extended Data Figure 8). In all species, the vDL2d neurons bypass the mushroom body and project only to the lateral horn. Therefore, the number of neurons connecting the DL2d glomerulus to the mushroom body is the same in all species. However, collectively, the adDL2d neurons show larger bouton volume in *D. sechellia* than they do in the other species (bouton volume of all adDL2d neurons: *D. melanogaster*: 112.18 ± 12.41; *D. simulans*: 104.44 ± 25.93; *D. sechellia*: 348.29 ± 186.63; Table 1, Figure 3d). We found that the increased number of connections between adDL2d neurons and Kenyon cells in *D. sechellia* is due to more presynaptic boutons being formed per neuron (Figure 4b-c, Extended Data Table 3). We also found that individual adDL2d neurons show more complex branching patterns based on several quantifiable parameters (Figure 4c, Extended Data Table 3). We identified a similar phenotype for the projection neurons associated with the DP1l glomerulus: there are three DP1l projection neurons in *D. sechellia* (one neuron located in the lateral cluster, or lDP1l neuron, and two vDP1l neurons) but only two DP1l projection neurons in *D. melanogaster* (one lDP1l neuron and one vDP1l neuron) as it has been reported in the connectomes^22^ (Table 1, Extended Data Figure 9). As with the vDL2d neurons, the vDP1l neurons bypass the mushroom body and only the lDP1l neurons connect to Kenyon cells. We found that for lDP1l neurons the bouton volume is larger in *D. sechellia* than in *D. melanogaster* and *D. simulans* (bouton volume of the lDP1l neuron: *D. melanogaster*: 244.39 ± 69.00; *D. simulans*: 232.41 ± 100.73; *D. sechellia*: 388.91 ± 75.88).

The DC3 and VM5v glomeruli are less frequently connected to Kenyon cells in *D. sechellia* (Figure 2d, Extended Data Table 2). We identified two DC3 projection neurons in *D. sechellia* (two adDC3 neurons) but as many as three in *D. melanogaster* (three adDC3 neurons as reported in the connectomes^22^) and six in *D. simulans* (three adDC3 neurons and three vDC3 neurons) (Table 1, Figure 3b-c, Extended Data Figure 8). In *D. simulans*, the vDC3 neurons bypass the mushroom body and project only to the lateral horn. The volume of presynaptic boutons collectively formed by the adDC3 neurons in the mushroom body is significantly smaller in *D. sechellia* than in *D. melanogaster* and *D. simulans* (bouton volume of all adDC3 neurons: *D. melanogaster*: 370.15 ± 195.68; *D. simulans*: 364.06 ± 146.60; *D. sechellia*: 233.02 ± 83.80; Table 1, Figure 3d). However, individual adDC3 projection neurons are morphologically similar across species with no significant differences in branch length, number of forks or bouton number (bouton volume of individual adDC3 neuron: *D. melanogaster*: 184.15 ± 34.60; *D. simulans*: 169.01 ± 43.23; *D. sechellia*: 165.36 ± 25.56; Figure 4b-c, Extended Data Table 3). Thus, the decrease in the number of connections between adDC3 neurons and Kenyon cells in *D. sechellia* is due to a decrease in number of projection neurons innervating the DC3 glomerulus in that species. We identified a similar phenotype for the projection neurons associated with the VM5v glomerulus: there are two adVM5v projection neurons in *D. sechellia* but as many as three adVM5v projection neurons in *D. melanogaster*, as reported in the connectomes^22^, and three adVM5v projection neurons in *D. simulans* (Table 1, Figure 3b-c, Extended Data Figure 8). We found that, as for the adDC3 projection neurons, the adVM5v neurons form collectively a smaller bouton volume in *D. sechellia* but, individually, the VM5v neurons are similar across species (bouton volume of all VM5v neurons: *D. melanogaster*: 256.78 ± 104.09; *D. simulans*: 268.43 ± 132.67; *D. sechellia*: 140.14 ± 77.76; Table 1; bouton volume of individual VM5v neurons: *D. melanogaster*: 116.45 ± 18.68; *D. simulans*: 85.38 ± 17.48; *D. sechellia*: 169.73 ± 39.67; Figure 4b-c, Extended Data Table 3). Thus, as with the adDC3 neurons, there are fewer VM5v neurons in *D. sechellia*, and, therefore, fewer connections between the VM5v glomerulus and Kenyon cells. Combined with the observations made for the DL2d and DP1l projection neurons, these results suggest that increases in glomerular representation in *D. sechellia* occur through increases in bouton number, whereas decreases in glomerular representation occur through decreases in the number of projection neurons associated with different glomeruli.

We did not detect significant interspecific differences for the VC3 and VL2a projection neurons although we identified these glomeruli as differently represented in at least one of the species investigated (Table 1, Extended Data Figure 9). This result suggests that our photo-labeling and dye-labeling techniques might not be sensitive enough to detect all the morphological features — other than number of projection neurons and bouton volume — that underlie shifts in connectivity. Alternatively, this result could reflect the relatively low stringency of the statistical tests used to identify significant shifts in connectivity frequencies — we used a false discovery rate of 0.10 — and it could suggest that these data points might be false positives. Although our photo-labeling and dye-labeling techniques might not fully capture the extent of morphological differences that exist between the species investigated, the interspecific differences they successfully identified seem specific and restricted to a small number of projection neurons. We have performed similar morphological analyses on neurons innervating glomeruli that we identified as identically represented in all three species, namely the DA2, DM2 and DP1m glomeruli. We could not detect any noticeable differences in the axonal termini that these projection neurons extend in the mushroom body (Table 1, Extended Data Figure 9).

### Increased connectivity enhances learning performance

We tested whether the changes in glomerular representation we observed in the mushroom body of the species we investigated lead to measurable differences in learning performance by using a well-established behavioral paradigm^34^. In this paradigm, flies were conditioned to associate a stimulus — either air perfumed with an odor or mineral oil — with electric shocks, and their preference for the conditioned stimulus was subsequently tested in a T-maze (Extended Data Figure 10). In short, we measured the number of flies seeking out the conditioned stimulus over the number of flies seeking out the unconditioned stimulus to derive a Performance Index. We used either a protocol that included a single regimen of electric shocks (single training) or a protocol that included six spaced regimens of electric shocks (spaced training). In a first series of experiments, we tested odors known to activate the glomeruli whose representation shifts most drastically across species: the DC3 and DL2d glomeruli. Farnesol, a yeast odor, strongly and selectively activates the OR83c-expressing neurons in *D. melanogaster*^31^ (Extended Data Table 4). The OR83c-expressing neurons project to the DC3 glomerulus, and the adDC3 projection neurons connect to Kenyon cells at a much higher frequency in *D. melanogaster* and *D. simulans* than in *D. sechellia* (Figure 2c, Extended Data Table 4). We found that farnesol could not trigger strong learned responses in any of the species in either the single or spaced training protocol (Figure 5a-b, Extended Data Figure 11). Hexanoic acid, a noni odor, strongly and selectively activates the IR75b-expressing neurons in *D. sechellia* and weakly activates multiple types of olfactory sensory neuron in *D. melanogaster* ^20,32^ (Extended Data Table 4). The IR75b-expressing neurons project to the DL2d glomerulus, and the adDL2d projection neurons connect with Kenyon cells at a high frequency in *D. sechellia*; the projection neurons associated with the glomeruli activated by hexanoic acid collectively connect at a similarly high frequency in *D. melanogaster* and *D. simulans* (Figure 2c, Extended Data Table 4). We found that hexanoic acid can trigger learned responses in *D. melanogaster* and *D. simulans* when using either the single or spaced training protocol but not in *D. sechellia* where weak learned responses of much smaller amplitudes are only observed when using the spaced training protocol (Figure 5a-b, Extended Data Figure 11). These results suggest that activation of a single glomerulus is insufficient to elicit robust learned responses, and that the activation of multiple glomeruli that collectively connect to Kenyon cells at high frequencies might be required for learning.

**Figure 5.**
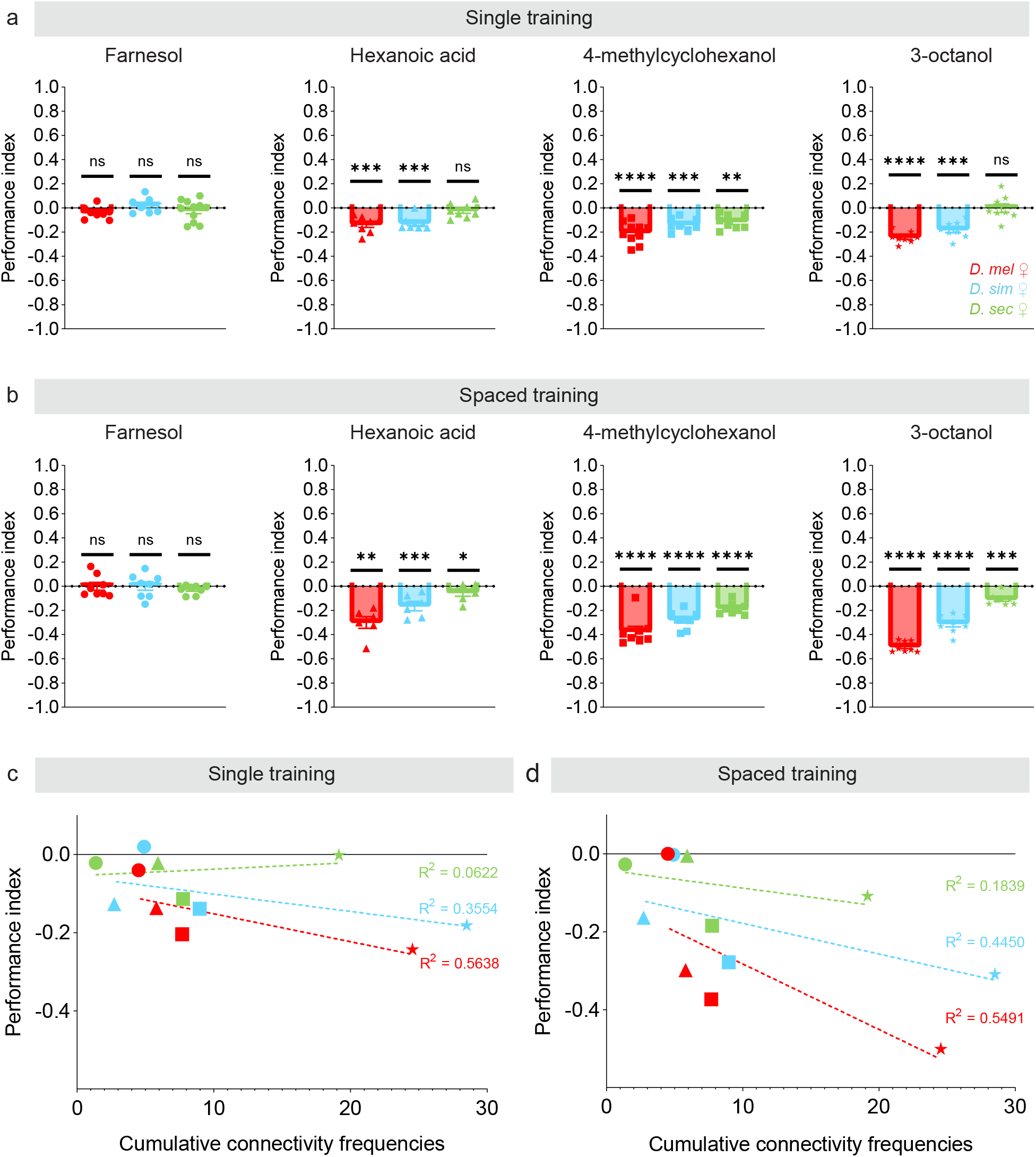
Learning abilities differ across species. (a,b) Flies (*D. melanogaster*: red; *D. simulans*: blue; *D. sechellia*: green) were trained to associate an odor (farnesol: circles, hexanoic acid: triangles, 4-methylcyclohexanol: squares or 3-octanol: stars) or its solvent (mineral oil) with punitive electric shocks using a single regimen of shocks (a) or six regimens of shocks (b) and learning was measured as a Performance Index (*n* ≥ 8); the Performance Indices obtained for the odor-pairing and the reciprocal pairing was averaged. See Extended Data Figure 12 for individual Performance Indices. The statistical significance, or ‘*p*-value’, was measured using the sample t test using ‘0’ as the hypothetical mean (*: *p*-value < 0.5, **: *p*-value < 0.01, ***: *p*-value < 0.001, ****: *p*-value < 0.0001). (c,d) The Performance Indices obtained for a given odor in a particular species (farnesol: circles, hexanoic acid: triangles, 4-methylcyclohexanol: squares or 3-octanol: stars) were plotted against the cumulative frequencies of the glomeruli known to be activated by a particular odor (based on a previous study^32^; Extended Data Table 4); the R^2^ values obtained for each regression line are shown.

We thus set out to test odors known to activate multiple types of olfactory sensory neuron in *D. melanogaster*, and therefore multiple glomeruli: 4-methylcyclohexanol and 3-octanol, which are detected by at least three and nine different types of olfactory sensory neuron, respectively^30,32^ (Extended Data Table 4). We found that both odors triggered strong learned responses in *D. melanogaster* using either the single or spaced training protocol (Figure 5a-b, Extended Data Figure 11). We noticed that the magnitude of the Performance Index varies significantly across odors: 3-octanol, which activates the largest number of glomeruli, gave rise to larger Performance Indices than 4-methycyclohexanol. To determine whether differences in connectivity frequencies could underlie the amplitude of the Performance Indices, we plotted the cumulative connectivity frequencies of the glomeruli known to be activated by each odor in *D. melanogaster* against the Performance Indices measured with a given protocol (Figure 5c-d). We observed a strong positive correlation, suggesting that odors activating a large number of highly connected glomeruli are more learnable than odors activating one or a few glomeruli. We performed similar analyses on *D. simulans* and *D. sechellia* and observed similar correlations. It is important to note that the cumulative connectivity frequencies were calculated using the odor responses measured in *D. melanogaster*, with the exception of hexanoic acid for which recordings collected in *D. simulans* and *D. sechellia* are available^20^. Interestingly, the Performance Indices also vary across species although all species display equally strong avoidance to electric shocks (Extended Data Figure 12): *D. melanogaster* performed better than *D. simulans* and remarkably better than *D. sechellia* regardless of the protocol used, whereas *D. sechellia* was only able to learn when we used the spaced training protocol (Figure 5c-d). These differences could reflect the fact that this training paradigm was originally developed and optimized for *D. melanogaster*; our data set indeed represents one of the first comparative study investigating differences in learning performance across *Drosophila* species. These differences could also reflect the fact that *D. sechellia* is impaired in its ability to synthesize dopamine, a neurotransmitter essential for learning^35^. The poor ability of *D. sechellia* to learn may reveal a relaxed requirement for this species to form associations with a wide range of stimuli as, unlike generalists, its survival depends on a single resource, noni. Regardless, our results clearly demonstrate that increased connectivity between projection neurons and Kenyon cells enhances learning performance.

## Discussion

In this study, we harness a phylogenetically and ecologically informed approach to pinpoint evolutionary differences in neuronal connectivity architecture and learning performance between three species of *Drosophila*. The species differ in the frequency with which inputs from a small set of olfactory channels — mainly those relaying information about food odors — are represented among the overall input to a higher-order processing center, the mushroom body. Notably, most of these differences are found in the specialist species *D. sechellia*, suggesting they are due to an ecological niche shift, rather than merely a function of phylogenetic distance. Evolutionary differences in sensory representation are caused by morphological alterations of the projection neurons connecting individual glomeruli to the mushroom body, either through changes in the number of projection neurons or through changes in the number of presynaptic boutons per neuron. Our study also shows that increased connectivity enhances learning performance in an associative task.

It is likely that other features of connectivity architecture underlie evolution of the mushroom body in *Drosophila* species. A recent electron microscopy-based study of the *D. melanogaster* mushroom body has revealed a subtle but significant connectivity pattern between a subgroup of projection neurons and the α/β and α’/β’ Kenyon cells, and our mapping technique may not be sensitive enough to capture these more subtle features of connectivity architecture^27,28^. Likewise, there are limitations regarding the forces driving the evolution of connectivity architecture in the mushroom body. For instance, it remains unclear whether the neuronal connectivity traits identified are variable in natural populations, especially those inhabiting in different environments. It remains equally unclear whether the anatomical and behavioral changes that we observed confer a fitness advantage and could be adaptive. The fact that we were able to correlate a visible neuronal phenotype with these connectivity changes and measure the impact of such changes on learning, make such investigations theoretically possible in the future.

Evolutionary comparisons of the olfactory system in *Drosophila* species have so far mainly focused on peripheral components of the circuit, in particular the olfactory sensory neurons and the receptor genes they express^36^. What has remained less clear is whether changes in receptor cell response profile or amplitude are sufficient to lead to evolutionary shifts in chemosensory ecology or whether concomitant changes occur in higher brain centers. Our study provides the first evidence at the cellular and behavioral levels for such evolutionary changes in the connectivity architecture of higher olfactory brain centers and how these changes might enhance cognitive functions. The fact that biases in connectivity are genetically encoded in *D. melanogaster*, and form independently of neuronal activity, indicates that the changes we uncovered are not merely a consequence of altered function at the periphery^28^. Instead, they represent an additional element of evolutionary change in circuit architecture and point to a locus in the central brain where evolution of neuronal circuits occurs.

How higher brain centers evolve remains, for the most part, completely unknown. Higher brain centers often feature more integrated circuits with many connections, and therefore it is conceivable that they might be less liable to change during evolution due to the disruptive impacts such changes might have across circuits. The anatomical changes we observed in this study are comparatively subtle, involving changes in the quantity of neuronal connections formed while leaving the overall random circuit architecture intact. Theoretical studies have demonstrated that randomization of input enables mushroom body-like networks to generate a representation space of high dimensionality, where many odors can be represented by non-overlapping neuronal ensembles^37–39^. We propose that biases in connectivity enable the mushroom body to better represent ecologically meaningful odors by increasing the coding space allocated to these odors, potentially increasing the fitness of an animal. Shifts in connectivity biases emphasize information streams that are most relevant to an animal without completely inactivating or adding *de novo* streams. This evolutionary pattern may be widespread across brain centers and species.

## ACKNOWLEDGMENTS

We thank members of the Caron laboratory for comments on the manuscript; David Stern for sharing the *attp^2176^* and *attp^2178^ D. simulans* strains and φC31 integrase mRNA; Vanessa Ruta for sharing the *UAS-C3PA-GFP* plasmid; Adam Lin for preparation of the standard cornmeal agar medium; the Cell Imaging Core at the University of Utah for use of the Zeiss LSM 880 microscope; Ashley Platt and Miles Jacob for assistance with general laboratory concerns. This work has been funded by grants from the National Institute for Neurological Disorders and Stroke (R01 EB 029858, R01 NS 106018 and R01 NS 1079790), the National Science Foundation (DBI 1707398 and IOS 2042397), the Gatsby Charitable Foundation, the European Research Council (European Research Council Advanced Grant 833548) and the Swiss National Science Foundation (Ambizione Grant PZ00P3 185743). Further financial support was provided by the Eunice Kennedy Shriver National Institute of Child Health and Human Development (T32-HD-007491) (K.E.E.), the Department of Energy Computational Science Graduate Fellowship (DE-SC0022158) (I.G.), the Burroughs Wellcome Foundation (A.L.K.), the McKnight Endowment Fund (A.L.K.), the Simons Collaboration on the Global Brain (A.L.K.), the Human Frontier Science Program (LT000461/2015-L) (T.O.A.) and the Georges S. and Dolores Eccles Foundation (S.J.C.C.).

## AUTHORS CONTRIBUTIONS

K.E.E. and S.J.C.C. conceived the project. H.M.S. generated and analyzed the antennal lobe reconstructions; K.E.E. photo-labeled projection neurons and Kenyon cells, dye-labeled individual projection neurons, and analyzed their morphology; K.E.E. generated the connectivity matrices; S.B. performed and analyzed all behavioral experiments; I.G. and A.L.K. performed the statistical analyses of the connectivity matrices; E.V. generated the similarity trees; T.O.A. and R.B. provided the *D. sechellia* transgenic lines and plasmids used to generate the *D. simulans* transgenes. K.E.E and S.J.C.C. wrote the manuscript with input from all other authors.

---

**Extended Data Table 1.** Comparison of glomerular volumes across species. Individual glomeruli were reconstructed and their volumes were measured in *D. melanogaster*, *D. simulans* and *D. sechellia* (*n* = 3 for each species, standard deviation is shown). The statistical significance, or ‘*p*-value’, was measured when comparing glomerular volumes across species using the Mann-Whitney U test. See Extended Data Figure 1 for antennal lobe reconstructions.

| Extended Data Table 1 \| Comparison of glomerular volumes across species. |  |  |  |  |  |  |
| --- | --- | --- | --- | --- | --- | --- |
| Glomerulus | Mean ± s.d. volume ($\mu\text{m}^3$) | | | *p*-values | | |
|  | *D. melanogaster* | *D. simulans* | *D. sechellia* | *Dmel vs Dsim* | *Dsim vs Dsec* | *Dmel vs Dsec* |
| D | 2464 ± 40 | 1479 ± 186 | 2405 ± 131 | 0.1717 | 0.166161 | 0.72114 |
| DA1 | 4391 ± 123 | 3379 ± 218 | 2766 ± 238 | 0.1717 | 0.166161 | 0.117068 |
| DA2 | 1147 ± 62 | 1276 ± 200 | 1196 ± 123 | 0.751188 | >0.999999 | >0.999999 |
| DA3 | 476 ± 10 | 455 ± 53 | 451 ± 56 | 0.751188 | >0.999999 | 0.72114 |
| DA4l | 1133 ± 182 | 916 ± 71 | 837 ± 52 | 0.542211 | 0.490571 | 0.117068 |
| DA4m | 1097 ± 4668 | 1035 ± 110 | 814 ± 86 | 0.751188 | 0.303 | 0.228933 |
| DC1 | 2530 ± 144 | 1433 ± 217 | 1249 ± 148 | 0.1717 | 0.490571 | 0.117068 |
| DC2 | 1487 ± 178 | 1380 ± 152 | 904 ± 62 | 0.751188 | 0.166161 | 0.117068 |
| DC3 | 1665 ± 172 | 1749 ± 201 | 1163 ± 89 | 0.751188 | 0.166161 | 0.117068 |
| DC4 | 1702 ± 147 | 1270 ± 116 | 1050 ± 26 | 0.1717 | 0.166161 | 0.117068 |
| DL1 | 2006 ± 232 | 1618 ± 297 | 1448 ± 103 | 0.542211 | 0.819477 | 0.117068 |
| DL2d | 1565 ± 334 | 2007 ± 185 | 3066 ± 296 | 0.303 | 0.166161 | 0.117068 |
| DL2v | 1712 ± 164 | 1381 ± 141 | 1798 ± 59 | 0.1717 | 0.166161 | 0.72114 |
| DL3 | 1531 ± 213 | 1561 ± 158 | 931 ± 81 | >0.999999 | 0.166161 | 0.117068 |
| DL4 | 687 ± 47 | 1027 ± 96 | 1010 ± 90 | 0.1717 | >0.999999 | 0.117068 |
| DL5 | 1614 ± 50 | 1765 ± 195 | 1394 ± 13 | 0.751188 | 0.166161 | 0.117068 |
| DM1 | 3408 ± 168 | 4238 ± 90 | 2057 ± 73 | 0.1717 | 0.166161 | 0.117068 |
| DM2 | 3139 ± 227 | 3169 ± 227 | 4594 ± 127 | 0.751188 | 0.166161 | 0.117068 |
| DM3 | 1846 ± 93 | 1542 ± 113 | 1053 ± 56 | 0.1717 | 0.166161 | 0.117068 |
| DM4 | 3341 ± 64 | 2399 ± 130 | 724 ± 40 | 0.1717 | 0.166161 | 0.117068 |
| DM5 | 1784 ± 194 | 1900 ± 164 | 968 ± 96 | 0.751188 | 0.166161 | 0.117068 |
| DM6 | 1854 ± 42 | 1632 ± 134 | 2002 ± 137 | 0.303 | 0.166161 | 0.117068 |
| DP1l | 3269 ± 172 | 3118 ± 211 | 2173 ± 68 | 0.751188 | 0.166161 | 0.117068 |
| DP1m | 5117 ± 47 | 3399 ± 164 | 2465 ± 241 | 0.1717 | 0.166161 | 0.117068 |
| V | 3837 ± 194 | 3429 ± 126 | 2200 ± 146 | 0.1717 | 0.166161 | 0.117068 |
| VA1d | 3785 ± 148 | 1668 ± 126 | 2157 ± 194 | 0.1717 | 0.166161 | 0.117068 |
| VA1v | 5275 ± 160 | 3048 ± 330 | 2676 ± 187 | 0.1717 | 0.490571 | 0.117068 |
| VA2 | 4204 ± 199 | 3474 ± 130 | 1987 ± 125 | 0.1717 | 0.166161 | 0.117068 |
| VA3 | 1937 ± 120 | 1627 ± 233 | 894 ± 50 | 0.303 | 0.166161 | 0.117068 |
| VA4 | 1894 ± 174 | 1244 ± 217 | 871 ± 124 | 0.1717 | 0.303 | 0.117068 |
| VA5 | 1606 ± 220 | 1400 ± 170 | 811 ± 87 | 0.542211 | 0.166161 | 0.117068 |
| VA6 | 2604 ± 38 | 1929 ± 107 | 1505 ± 115 | 0.1717 | 0.166161 | 0.117068 |
| VA7l | 1388 ± 80 | 1285 ± 291 | 963 ± 98 | >0.999999 | 0.490571 | 0.117068 |
| VA7m | 1554 ± 144 | 783 ± 81 | 791 ± 105 | 0.1717 | >0.999999 | 0.117068 |
| VC1 | 1495 ± 214 | 1073 ± 66 | 816 ± 99 | 0.1717 | 0.166161 | 0.117068 |
| VC2 | 1382 ± 97 | 1140 ± 176 | 918 ± 121 | 0.303 | 0.303 | 0.117068 |
| VC3l | 1911 ± 207 | 1912 ± 111 | 1265 ± 117 | >0.999999 | 0.166161 | 0.117068 |
| VC3m | 1644 ± 187 | 1856 ± 253 | 2236 ± 180 | 0.542211 | 0.490571 | 0.117068 |
| VC4 | 1456 ± 159 | 1053 ± 152 | 844 ± 105 | 0.1717 | 0.490571 | 0.117068 |
| VC5 | 1408 ± 285 | 1238 ± 65 | 1294 ± 172 | 0.751188 | >0.999999 | 0.72114 |
| VL1 | 4319 ± 169 | 3062 ± 249 | 2503 ± 83 | 0.1717 | 0.166161 | 0.117068 |
| VL2a | 3222 ± 184 | 2014 ± 186 | 1954 ± 88 | 0.1717 | >0.999999 | 0.117068 |
| VL2p | 2912 ± 278 | 1983 ± 180 | 2137 ± 181 | 0.1717 | 0.819477 | 0.117068 |
| VM1 | 1397 ± 44 | 961 ± 93 | 1095 ± 109 | 0.1717 | 0.490571 | 0.117068 |
| VM2 | 1505 ± 51 | 1869 ± 104 | 1225 ± 116 | 0.1717 | 0.166161 | 0.117068 |
| VM3 | 1689 ± 135 | 2225 ± 193 | 1157 ± 114 | 0.1717 | 0.166161 | 0.117068 |
| VM4 | 1862 ± 131 | 1476 ± 29 | 822 ± 57 | 0.1717 | 0.166161 | 0.117068 |
| VM5d | 905 ± 115 | 1773 ± 127 | 3793 ± 153 | 0.1717 | 0.166161 | 0.117068 |
| VM5v | 885 ± 112 | 1311 ± 140 | 1258 ± 75 | 0.1717 | >0.999999 | 0.117068 |
| VM7d | 2006 ± 128 | 1139 ± 148 | 983 ± 138 | 0.1717 | 0.490571 | 0.117068 |
| VM7v | 2074 ± 65 | 1114 ± 123 | 766 ± 101 | 0.1717 | 0.166161 | 0.117068 |

**Extended Data Table 2.**
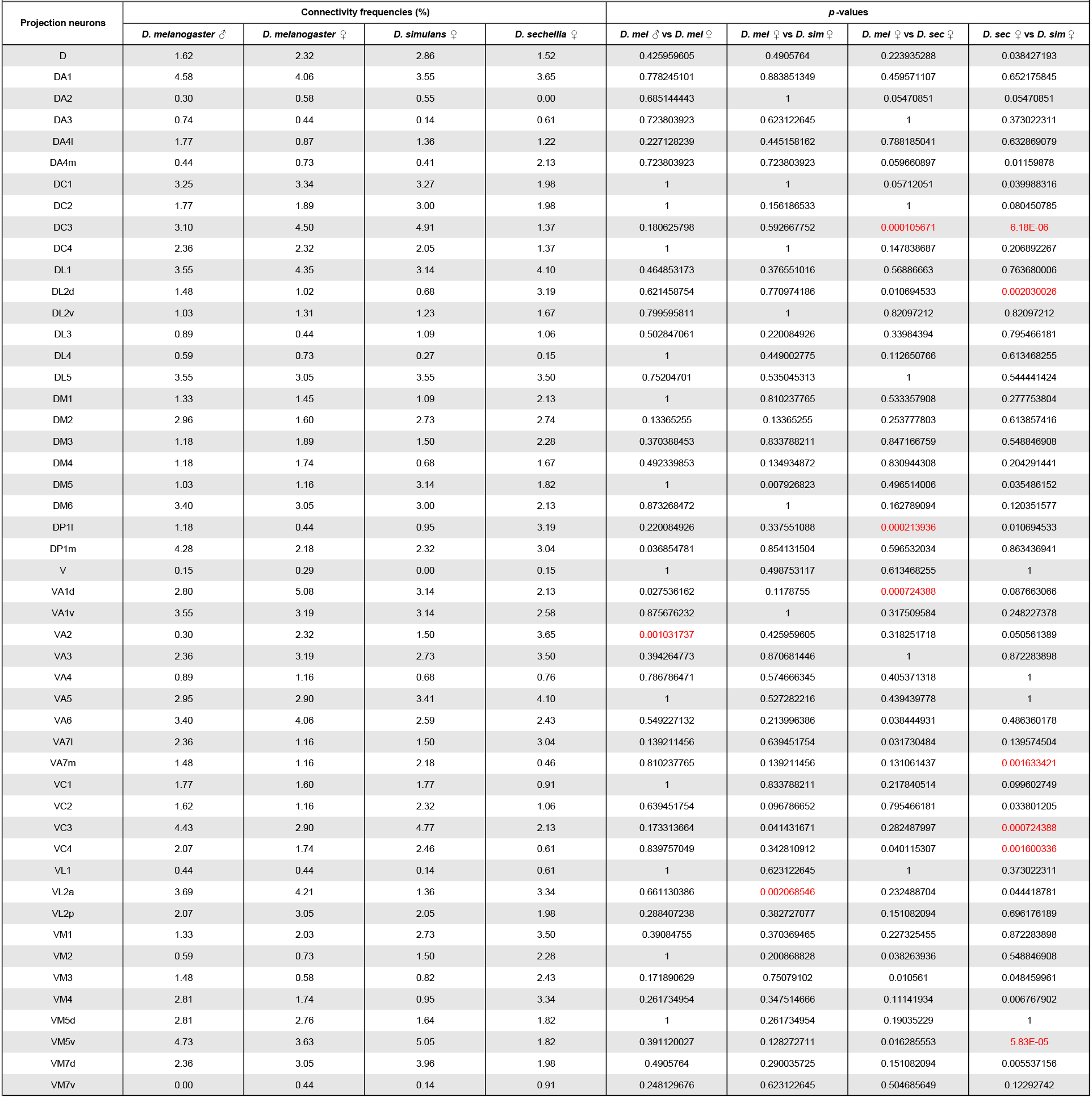
Comparison of connectivity frequencies across species. The frequencies at which projection neurons innervating a given glomerulus are connected to Kenyon cells were measured based on the number of connections detected in the connectivity matrices shown in Figure 1d. The statistical significance, or ‘*p*-value’, was measured when comparing connection frequencies across species using a binomial test; red text indicates *p*-value < 0.01.

**Extended Data Table 3.**
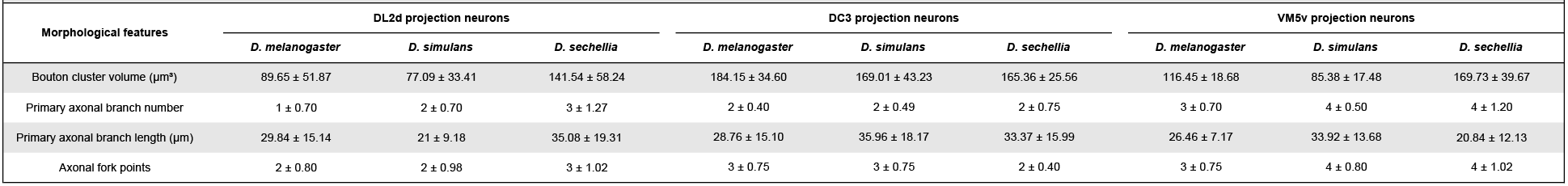
Morphological features of dye-labeled projection neurons across species. Morphological features of individual DC3, DL2d and VM5v projection neurons — namely the presynaptic bouton volume, the number of axonal primary branches and their length, and the number of forks they form in the mushroom body — were measured and compared across species (*n* = 5 for DC3 neurons, *n* = 10 for DL2d neurons and *n* = 5 for VM5d neurons, standard deviation is shown). See Figure 4b for representative images of each projection neuron type.

**Extended Data Table 4.** Connectivity frequencies and performance indices. Based on previous studies^20,32^, glomeruli were determined to be activated by a particular odor if that odor can elicit at least 0.25 of the maximal possible response in the olfactory sensory neurons associated with that glomerulus. Cumulative frequencies were calculated by adding the connectivity frequencies measured for each of the glomeruli activated by a given odor in each species.

| Extended Data Table 4 |  |  |  |  |
| --- | --- | --- | --- | --- |
| Odors used | Glomeruli activated | Cumulative connectivity frequencies (%) |  |  |
|  |  | *D. melanogaster* | *D. simulans* | *D. sechellia* |
| Farnesol | DC3 | 4.5 | 4.91 | 1.37 |
| Hexanoic acid | DL2d (*D.sec*), DM2 (*D.mel*, *D.sim*, *D.sec*), VL2a (*D.mel*) | 5.81 | 2.73 | 5.93 |
| 4-methylcyclohexanol | D, DA2, DM2, VA3 | 7.69 | 8.97 | 7.76 |
| 3-octanol | D, DC1, DC2, DM2, DM3, DM6, VA4, VC3, VM5d, VM5v | 24.54 | 28.5 | 19.16 |

**Extended Data Figure 1.**
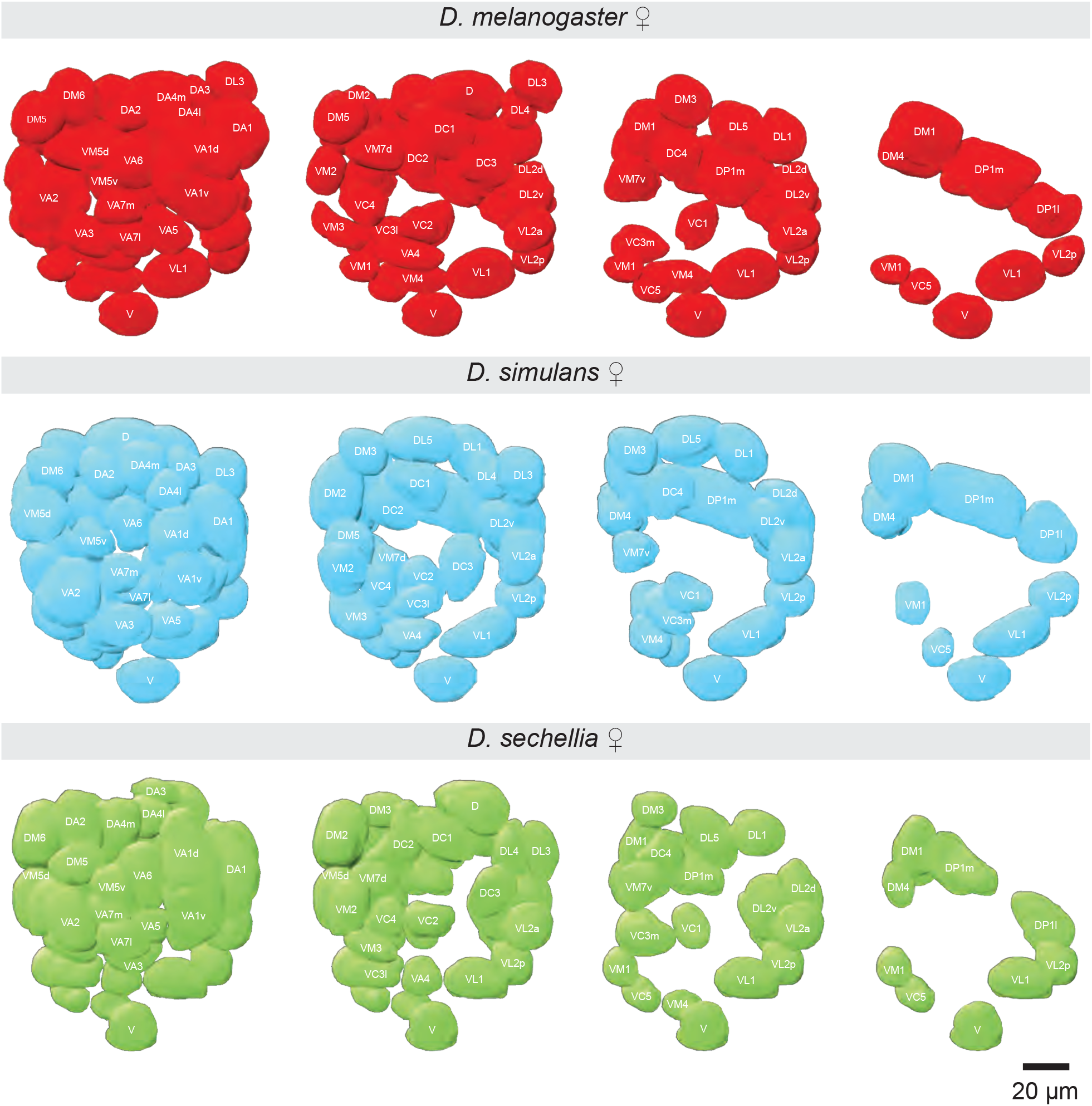
Antennal lobe reconstructions in different species. The brains of two-or three-day old *D. melanogaster* (red), *D. simulans* (blue), and *D. sechellia* (green) female flies were fixed, immuno-stained (using the nc82 monoclonal antibody against Bruchpilot) and imaged. The glomeruli forming the antennal lobe were individually reconstructed and identified based on shape and location. Three different planes are shown for each reconstruction. Scale bar is 20 μm. See Extended Data Table 1 for quantifications.

**Extended Data Figure 2.**
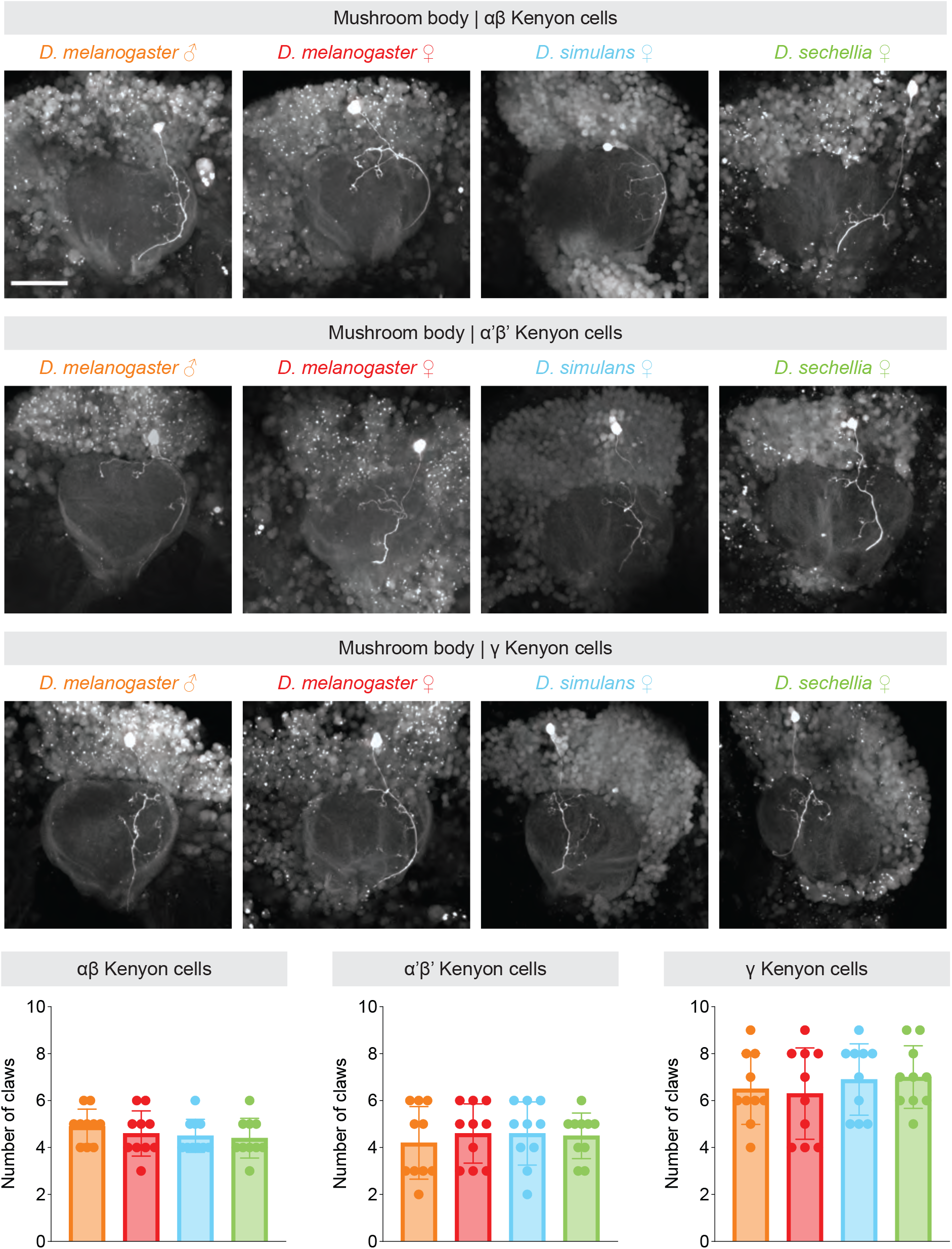
Morphological features of Kenyon cells across species. Individual. α/β (top panels), α’/β’ (middle panels), and γ Kenyon cells (bottom panels) were photo-labeled in *D. melanogaster* males (first column), *D. melanogaster* females (second column) *D. simulans* (third column), and *D. sechellia* flies (fourth column), and the postsynaptic terminals formed by these neurons in the mushroom body calyx — called ‘claws’ — were imaged. Scale bar is 50 μm. The total number of claws per Kenyon cell were counted for different types of Kenyon cell (orange: *D. melanogaster* males; red: *D. melanogaster* females; blue: *D. simulans*; green: *D. sechellia*; *n* = 10, standard deviation is shown). The statistical significance, or ‘*p*-value’, was measured to compare the number of claws measured for a given type of Kenyon cell using the Mann-Whitney U test but none of the values were found to be significantly different.

**Extended Data Figure 3.**
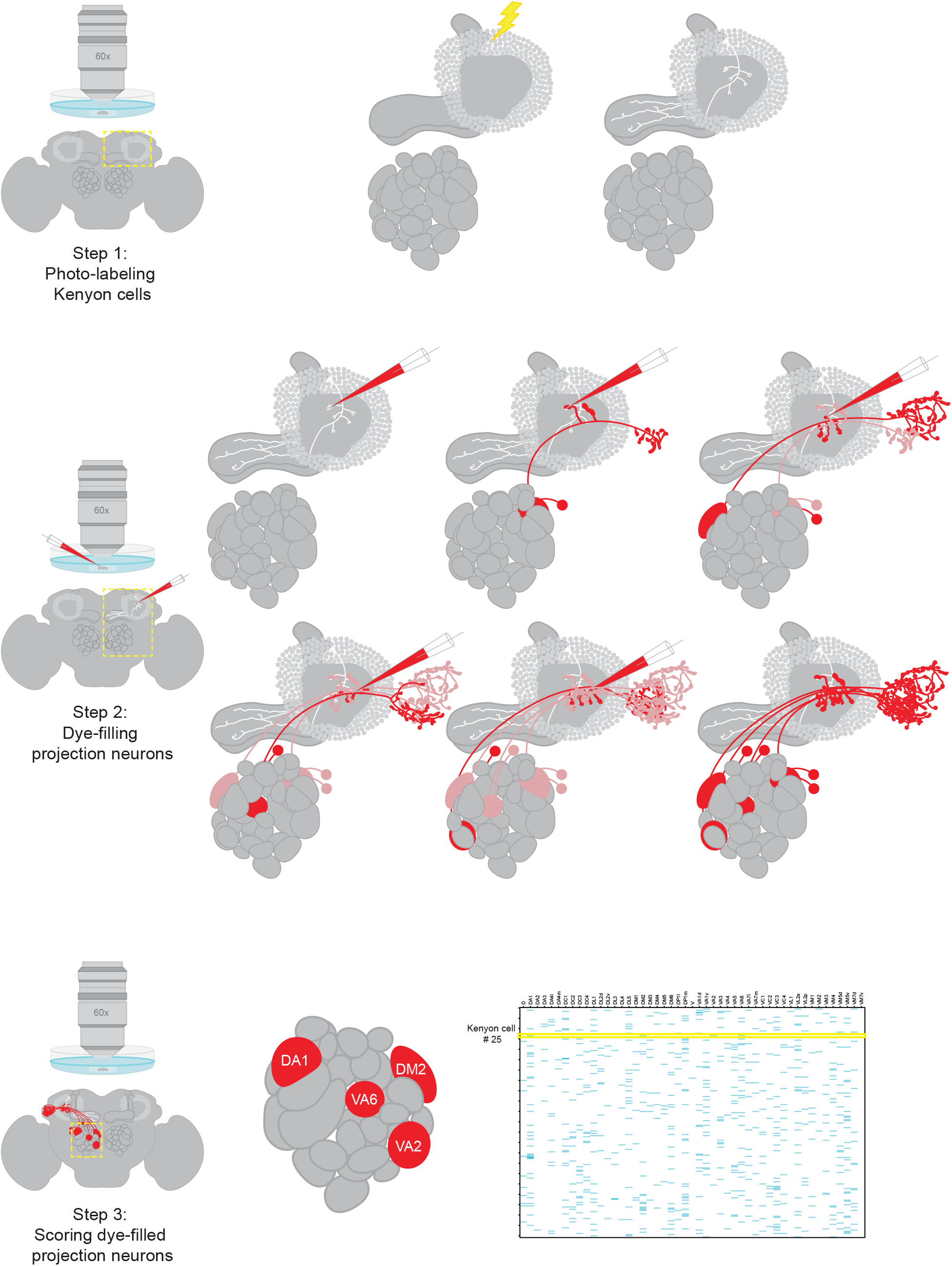
Mapping technique. Schematic depicting the three-step technique used to to map connections between projection neurons and Kenyon cells. Step 1 — Photo-labeling Kenyon cells: The brain of a fly carrying the *nSynaptobrevin-GAL4* and *UAS-photoactivatable-GFP* transgenes is dissected and imaged using 2-photon microscopy (yellow dashed box); a randomly selected Kenyon cell is targeted with high energy light (yellow lightning bolt) such that the photoactivatable-GFP molecules is converted only in that cell; the converted photoactivatable-GFP molecules rapidly diffuse in the Kenyon cell revealing its entire morphology (white). Step 2 — Dye-filling projection neurons: A post-synaptic terminal — or ‘claw’ — is targeted with an electrode filled with Texas Red dextran dye (red), and, following a short current pulse, the dye is electroporated into the projection neuron connected to that claw; this procedure can be repeated with other claws. Step 3 — Scoring dye-filled projection neurons: The identity of the dye-labeled projection neurons connected to the photo-labeled Kenyon cell can be revealed by visualizing the antennal lobe (yellow dashed box); the glomerular inputs of a given Kenyon cell — in this example the DA1, DM2, VA2 and VA6 glomeruli connecting to Kenyon cell #25 — is reported as a line in a connectivity matrix; each matrix reports the inputs identified for 200 Kenyon cells.

**Extended Data Figure 4.**
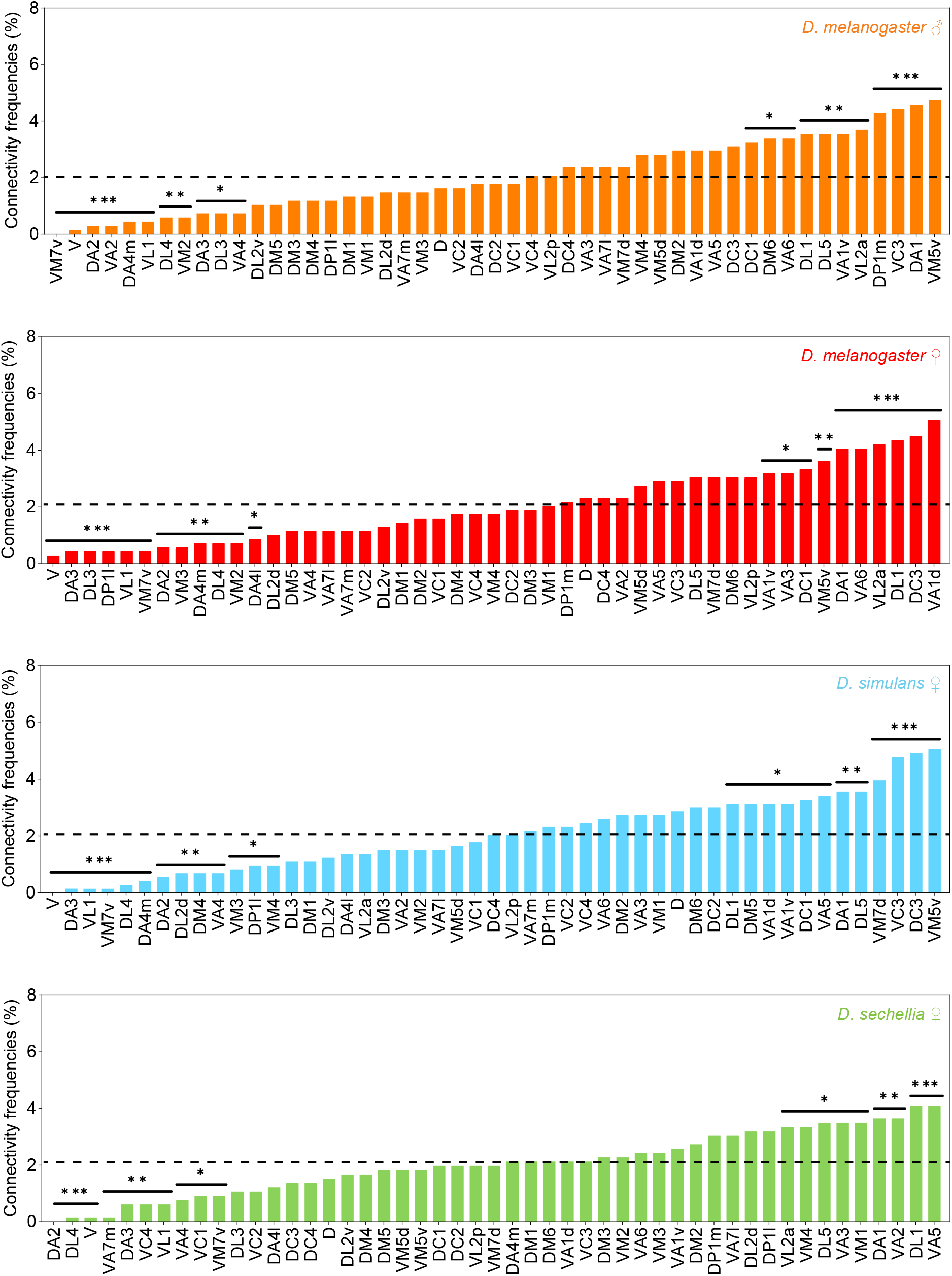
Non-uniform distribution of connectivity frequencies. The frequencies at which individual glomeruli are connected to Kenyon cells was determined based on the number of connections detected between projection neurons and Kenyon cells in each connectivity matrix reported in Figure 1d (orange: *D. melanogaster* males; red: *D. melanogaster* females; blue: *D. simulans*; green: *D. sechellia*). Glomeruli that are significantly underrepresented or overrepresented are labeled with asterisks (*: *p*-value < 0.5, **: *p*-value < 0.01, ***: *p*-value < 0.001).

**Extended Data Figure 5.**
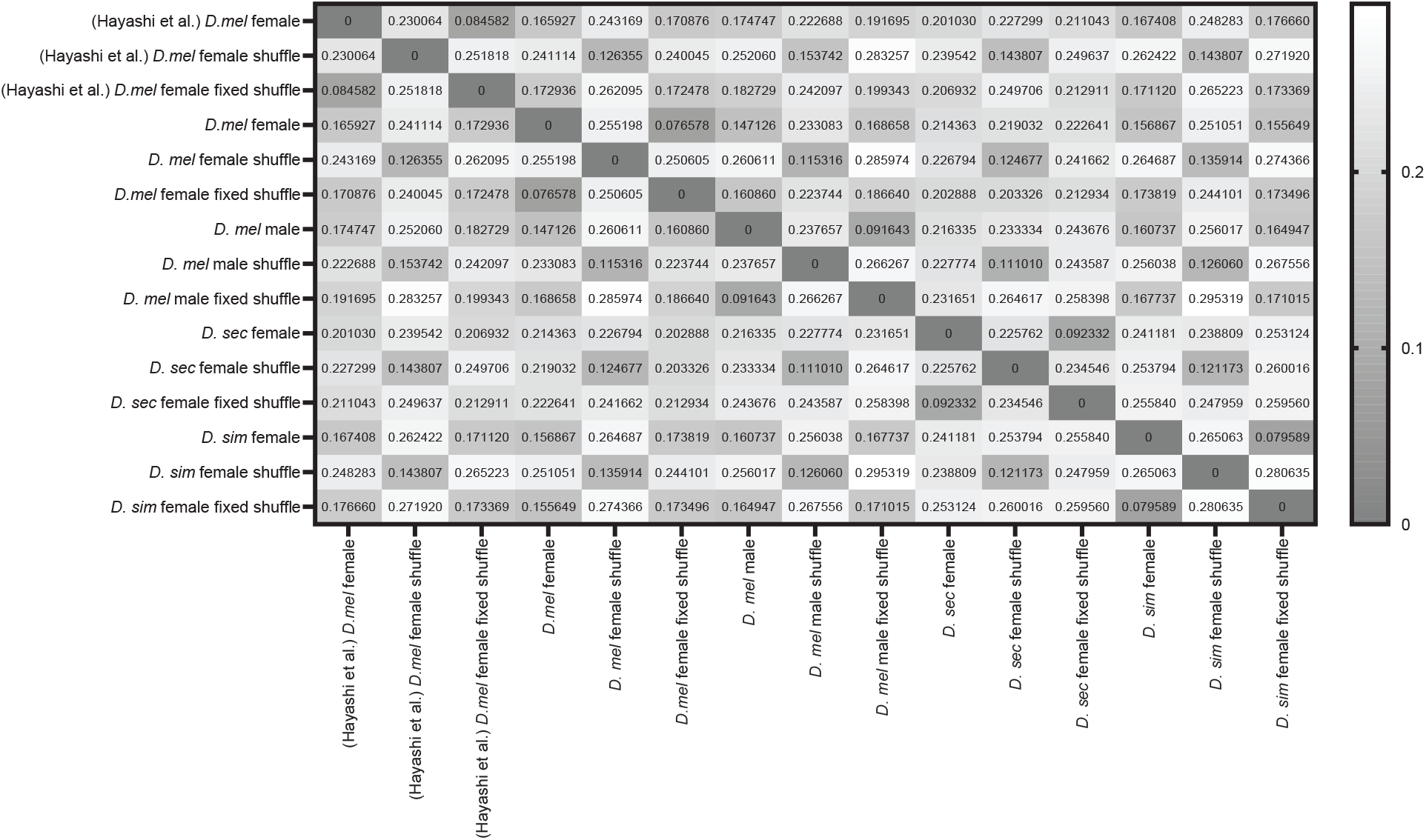
Jensen-Shannon distances. The Jensen-Shannon distances were measured by comparing the distributions in connectivity frequencies observed in the experimental matrices reported in this study(Figure 1d) or in a previous study^28^ and the uniform shuffle and biased shuffle matrices; distances are reported as a heat map. The color bar denotes the length of the distances measured.

**Extended Data Figure 6.**
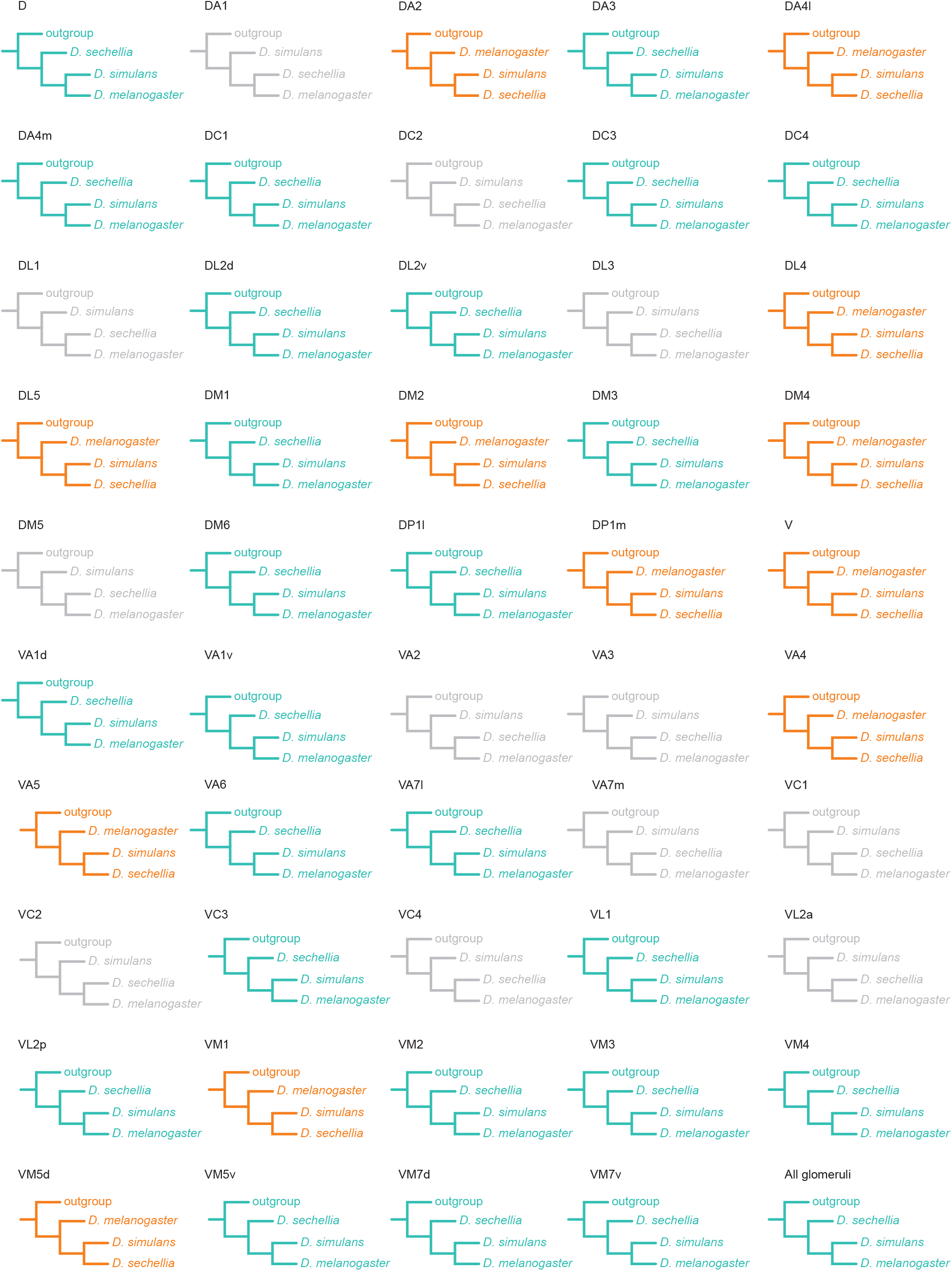
Similarity trees built using connectivity frequencies. Similarity trees were generated with the Tree analysis Using New Technology (TNT) software. The connectivity frequencies measured for a given glomerulus in the different species investigated were used as traits; a value of 100/49 or 2.04816 was used for the glomeruli forming the outgroup. Trees were generated either for individual glomeruli or for all glomeruli; trees either follow the known phylogenetic relationships (orange), the ecological relationships (aqua) or neither (grey).

**Extended Data Figure 7.**
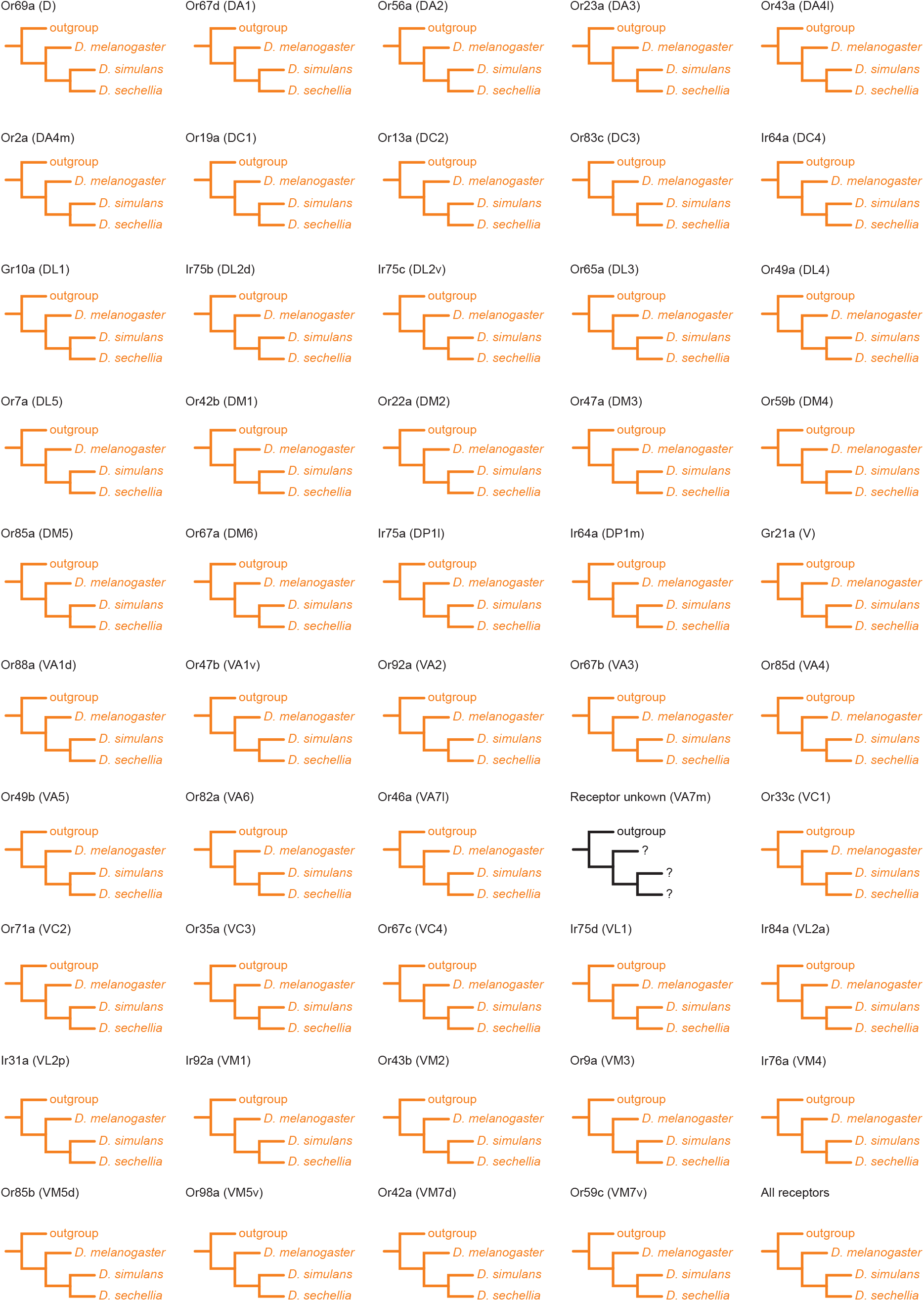
Similarity trees built using receptor sequences. Similarity trees were generated with the Tree analysis Using New Technology (TNT) software. The cDNA sequence of the receptor gene expressed by the olfactory sensory neurons associated with a given glomerulus, if known, was used as traits; the cDNA sequences of *D. yakuba* (Gr10a, Ir31a, Ir76a, Or9a, Or22a, Or23a, Or46a, Or49a, Or65a, OR67a, Or67d, Or69a, Or82a, Or85a, Or85b, Or85d, Or88a, Or92a, Or98a) or *D. elegans* (Gr21a, Ir64a, Ir75a, Ir75b, Ir75c, Ir75d, Ir84a, Ir92a, Or2a, Or7a, Or13a, Or19a, Or33c, Or35a, Or42a, Or42b, Or43a, Or43b, Or47a, Or47b, Or49b, Or56a, Or59b, Or59c, Or67b, Or67c, Or71a, Or83c) were used as the outgroup. Trees were generated either for individual glomeruli or for all glomeruli; all trees follow the known phylogenetic relationships (orange).

**Extended Data Figure 8.**
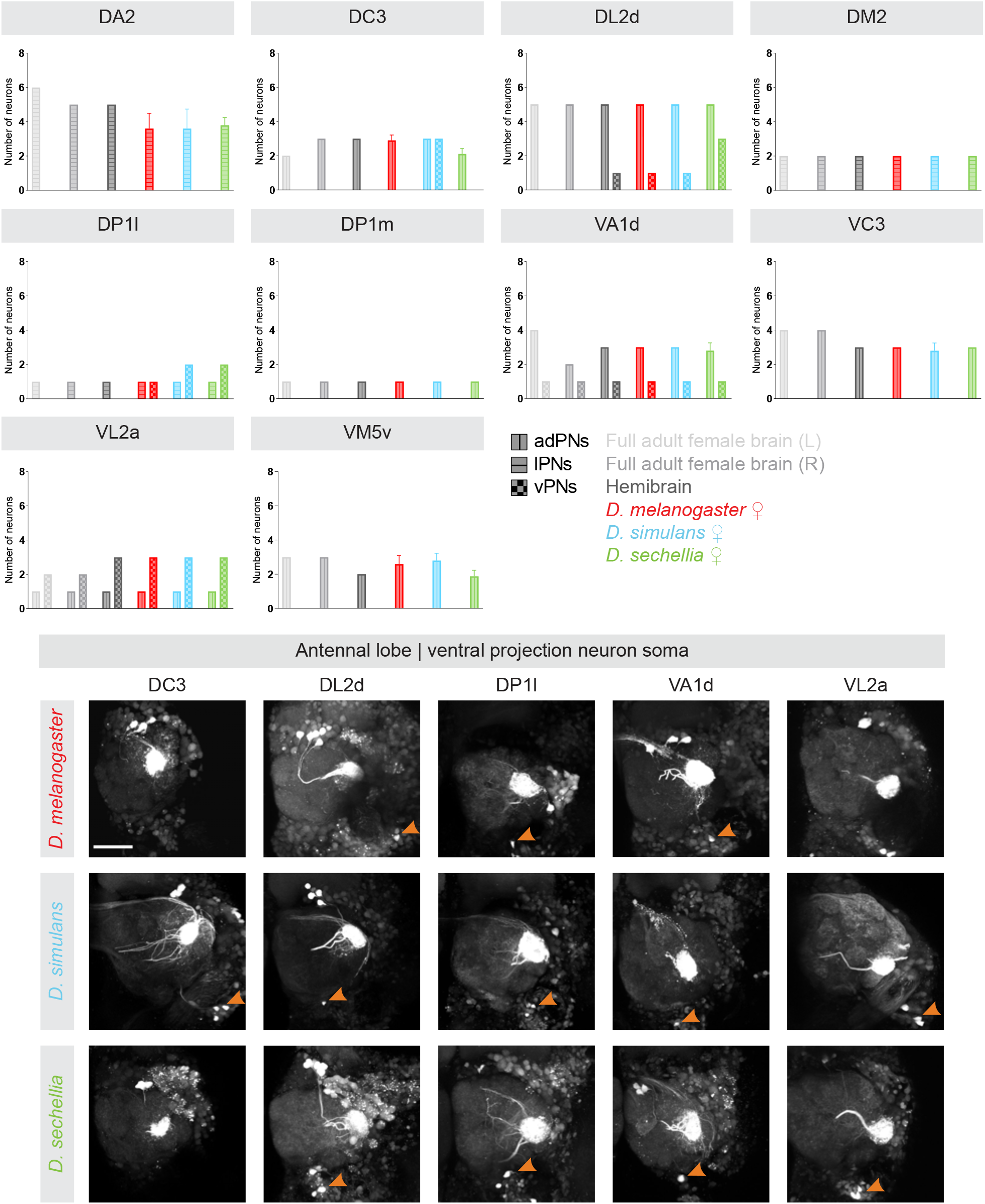
Number of projection neurons per glomerulus. The number of projection neurons associated with the DA2, DC3, DL2d, DM2, DP1l, DP1m, VA1d, VC3, VL2a, and VM5v glomeruli — divided by types (vertical lines: projection neurons from the anterior-dorsal clusters or adPNs; horizontal lines: projection neurons from the lateral clusters or lPNS; checkers: projection neurons from the ventral clusters or vPNs) — were compared across species based on the data sets collected in this study and the available *D. melanogaster* connectomes^22^ (light grey: FAFB connectome; medium grey: FAFB connectome; dark grey: hemibrain connectome). The cell bodies of the DC3, DL2d, DP1l, VA1d, and VL2a projection neurons that are located in the ventral cluster (orange arrows) that were photo-labeled in *D. melanogaster* (upper panels), *D. simulans* (middle panels) and *D. sechellia* (lower panels) are shown. Scale bar is 50 µm.

**Extended Data Figure 9.**
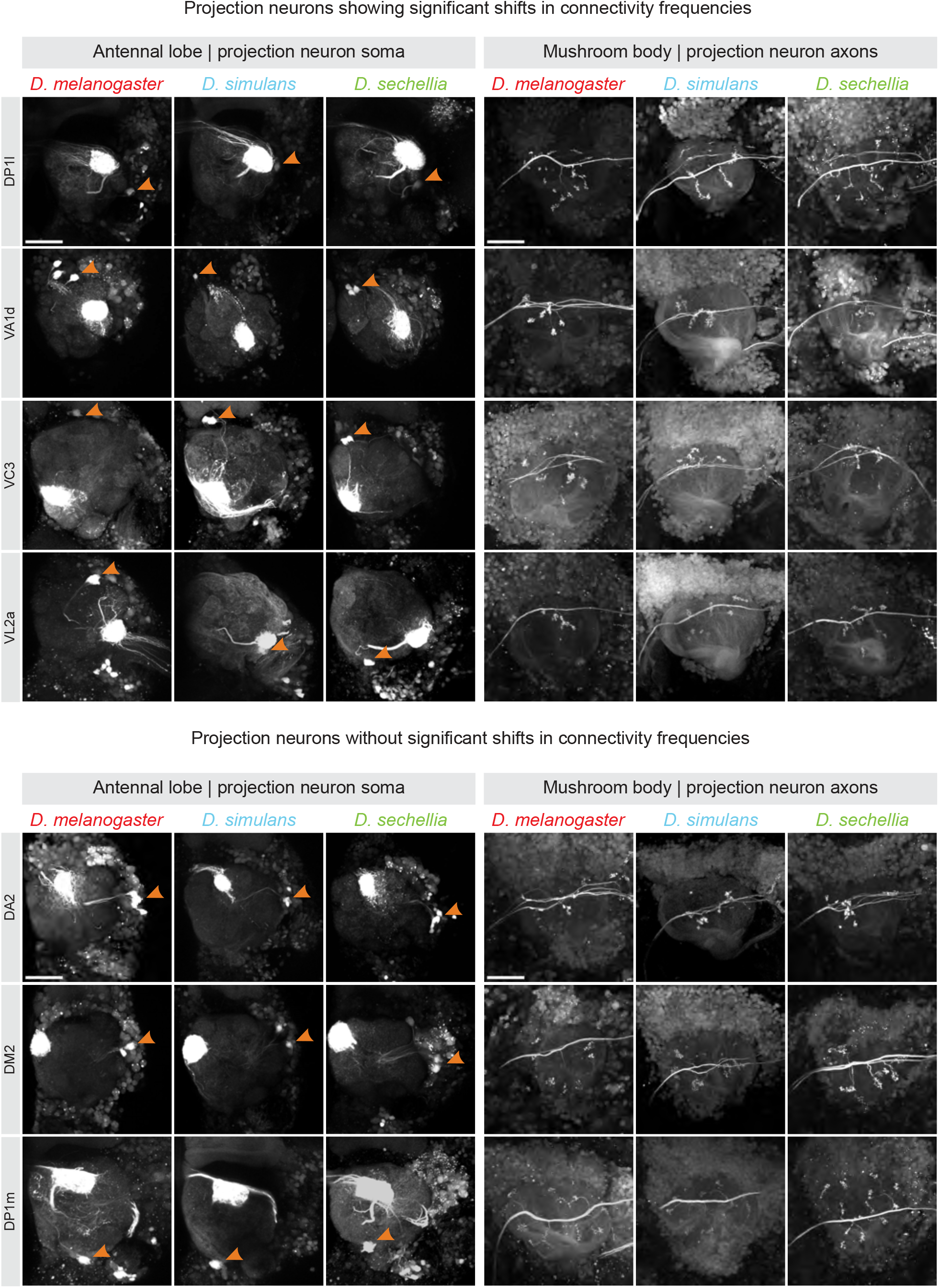
Morphological features of projection neurons across species. Projection neurons showing significant shifts in connectivity frequencies (DP1l, VA1d, VC3 and VL2a) were analyzed as well as projection neurons showing no such shifts (DA2, DM2 and DP1m). Projection neurons were photo-labeled in *D. melanogaster*, *D. simulans* and *D. sechellia*; the location of the cell bodies of the photo-labeled neurons relative to the antennal lobe (orange arrows) and the morphology of the axonal termini that these neurons extend in the mushroom body were imaged. Scale bar is 50 µm. See Table 1 for quantifications.

**Extended Data Figure 10.**
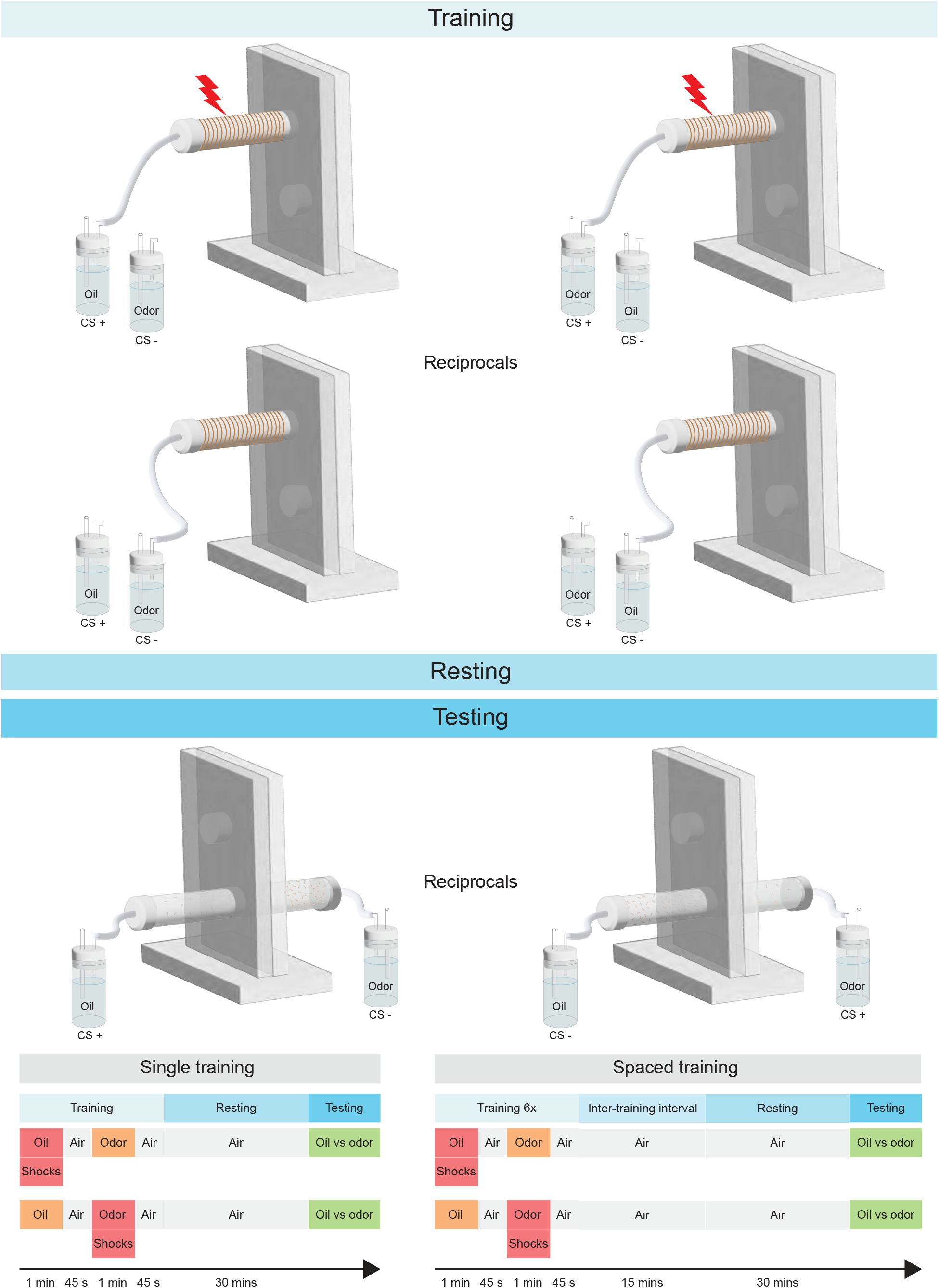
Aversive learning paradigm. Flies were trained in an aversive learning paradigm using two different protocols. During the training phase, flies were presented with either an odor (farnesol, hexanoic acid, 4-methylcyclohexanol or 3-octanol) or mineral oil while receiving electric shocks (CS+); soon after, flies were exposed to the other stimulus without experiencing electric shocks (CS-); in the reciprocal experiment, the reverse pairing was performed. Flies were allowed a resting phase of 30 minutes. During the testing phase, the preference of flies to seek out the CS+ over the CS-was measured and reported as a Performance Index. Flies were trained using a protocol that included a single regimen of twelve electric shocks or a protocol that included six spaced regimens of electric shocks.

**Extended Data Figure 11.**
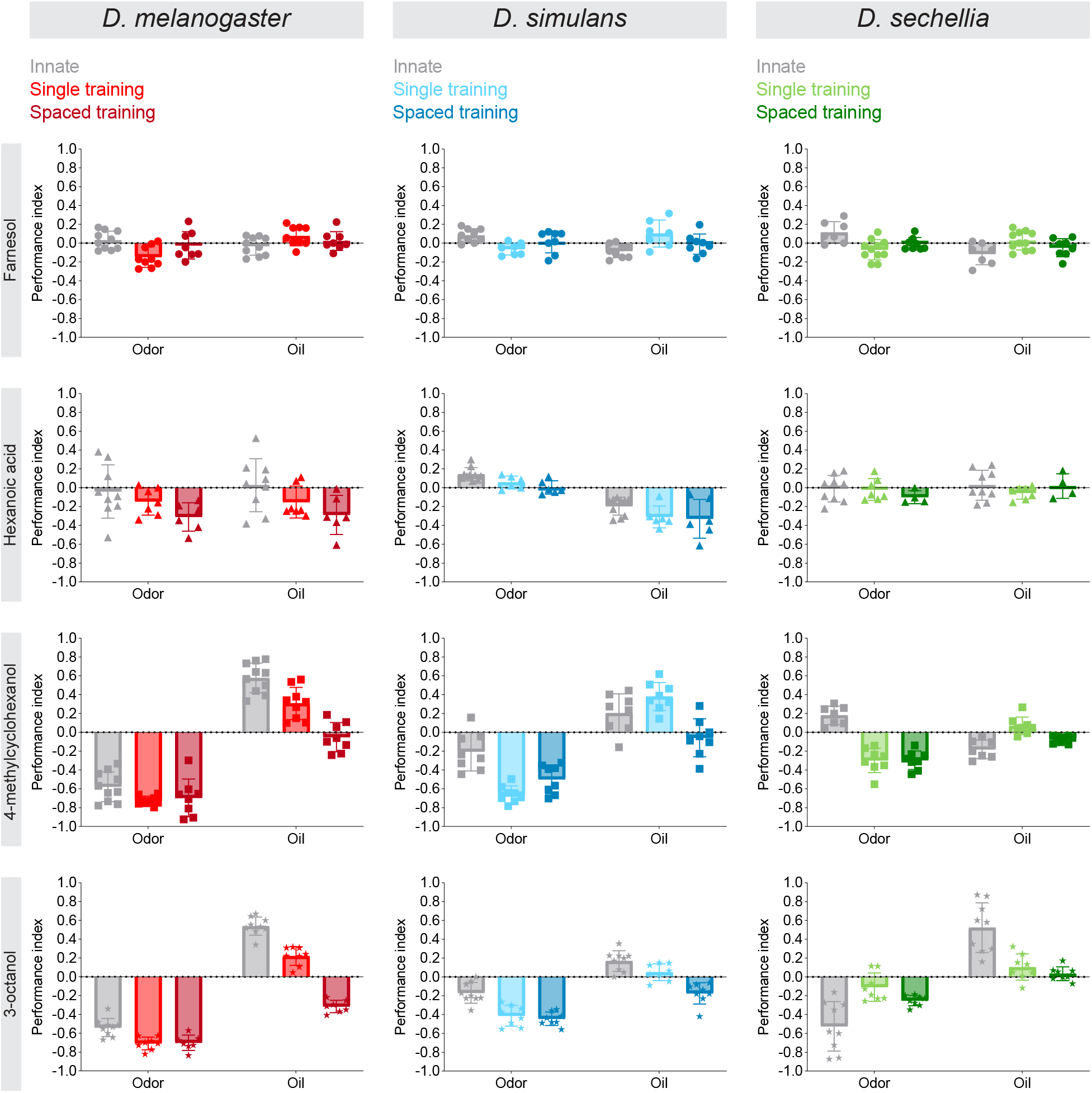
Single and spaced training. Flies (*D. melanogaster*: red and left column; *D. simulans*: blue and middle column; *D. sechellia*: green and right column) were trained to associate an odor (hexanoic acid: triangles, farnesol: circles, 4-methylcyclohexanol: squares or 3-octanol: stars) with punitive electric shocks using a single regimen of shocks (bright red, blue and green) or six regimens of shocks (dark red, blue and green) and learning was measured as a Performance Index (*n* ≥ 8); the response of flies to the stimulus before training is shown (gray). Plots showing the Performance Indices obtained for the odor-pairing (odor) and the reciprocal training (mineral oil) are shown.

**Extended Data Figure 12.**
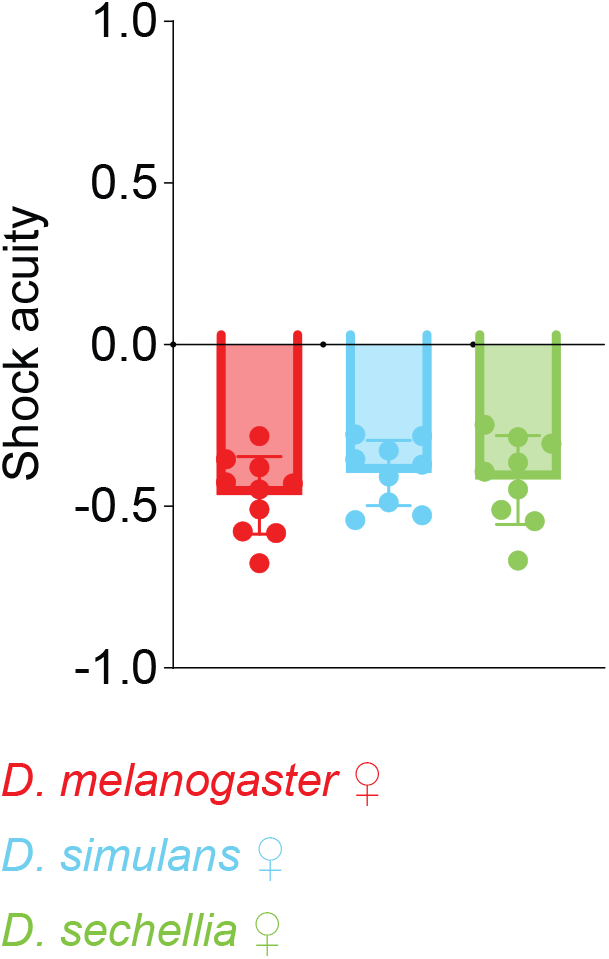
Shock acuity. Shock acuity was measured by allowing flies (*D. melanogaster*: red; *D. simulans*: blue; *D. sechellia*: green) to choose between a chamber lined with a copper grid onto which 90V electric shocks were delivered every five seconds over the course of one minute. The statistical significance, or ‘*p*-value’, was measured using the Anova test but none of the values were found to be significantly different.

## METHODS

### Fly stocks and husbandry

Flies were reared under standard conditions (25°C, 60% humidity) in incubators that maintain a 12h light/12h dark cycle (Percival Scientific Inc, Cat#DR36VL); *D. melanogaster* and *D. simulans* flies were reared on standard cornmeal agar medium, whereas *D. sechellia* flies were reared on standard cornmeal agar medium that was supplemented with noni juice (Healing Noni). The stocks used and their sources were as follows: *D. melanogaster*: *w^1118^;;;* (Bloomington Stock Center, 5905)*, yw;[N-Synaptobrevin-GAL4]^2.1^;;* (J. Simpson, University of California, Santa Barbara)^23^ and *y^1^,w^1118^;[10xUAS-IVS-Syn21-mC3PA-GFP-p10]*^attP40^;; (Axel laboratory, Columbia University)^23^; *D. sechellia*: *w[N-Synaptobrevin-GAL4, w*]3P3-RFP-DEL;;;* and *w[UAS-C3PA-GFP, w*]3P3-RFP-DELA;;;* (Benton laboratory, University of Lausanne)^16^; *D.* simulans: *attp^2176^* y*^1^w^1^; pBac [3P3-EYFP-DEL, attp];;* and *attp^2178^ y^1^w^1^;; pBac [3P3-EYFP-DEL, attp];* (Stern laboratory, Janelia Farm Research Campus)^40^.

### Reconstructing antennal lobes

Antennal lobes were reconstructed from confocal images of immuno-stained brains. The brains of flies were dissected at room temperature in a phosphate buffered saline solution or PBS (Sigma-Aldrich, P5493), fixed in 2% paraformaldehyde (Electron Microscopy Sciences, 15710) for either 45 min (*D. melanogaster*) or 35 min (*D. simulans* and *D. sechellia*) at room temperature, washed five times in PBST (PBS with 1x Triton, Sigma-Aldrich, T8787) at room temperature, blocked with 5% goat Serum (Jackson ImmunoResearch Laboratories) in PBST for 30 min at room temperature, and incubated in a solution that contained the primary antibody (1:20 in 5% Goat Serum/PBST, Developmental Studies Hybridoma Bank, nc82, AB 2314866) at 4°C overnight. On the following day, brains were washed four times in PBST and incubated in a solution that contained the secondary antibody (1:500 in 5% Goat Serum/PBST, Thermal Fisher, goat anti-mouse Alexa Fluor 488, AB 2576217) at 4°C overnight. On the following day, brains were washed four times in PBST and mounted on a slide (Fisher Scientific, 12-550-143) using the mounting media VECTASHIELD (Vector Laboratories Inc., H-1000). Immuno-stained brains were imaged using an LSM 880 confocal microscope (Zeiss). Each antennal lobe was reconstructed from a confocal image using the segmentation software Amira (FEI Visualization Sciences Group, version 2020.3.1). Individual glomeruli were reconstructed via manual segmentation: boundaries were demarcated by hand and interpolated. Glomeruli were assigned identities according to their position based on the available anatomical maps and the *D. melanogaster* hemibrain connectome v1.2.1^13,14,29,41^. Glomerular volumes were calculated from the reconstructed voxel size, and the sum of those volumes were used to calculate whole antennal lobe volumes. We identified a total of 51 glomeruli in the antennal lobe reconstructions but only 49 in the mapping experiments used to generate the connectivity matrices. This is because VC3 is split into two glomeruli — VC3m and VC3l — in the reconstructions but when scoring matrices, we could not distinguish VC3m from VC3l.

### Photo-labeling projection neurons and Kenyon cells

Neurons were photo-labeled based on a previously published protocol^42^. In short, brains were dissected in saline (108 mM NaCl, 5 mM KCl, 5 mM HEPES, 5 mM Trehalose, 10 mM Sucrose, 1 mM NaH_2_PO_4_, 4 mM NaHCO_3_, 2 mM CaCl_2_, 4 mM MgCl_2_, pH≈7.3), treated for 1 min with 2 mg/ml collagenase (Sigma-Aldrich) and mounted on a piece of SYLGARD placed at the bottom of a Petri dish. Each brain was either mounted with its anterior side facing upward (for photo-labeling projection neurons) or with its posterior side facing upward (for photo-labeling Kenyon cells). The photo-labeling and image acquisition steps were performed using a two-photon laser scanning microscope (Bruker, Ultima) with an ultrafast Chameleon Ti:Sapphire laser (Coherent) modulated by Pockels Cells (Conotopics). During the photo-labeling step, the laser was tuned to 710 nm and about 5 to 30 mW of laser power was used; during the image acquisition step, the laser was tuned to 925 nm and about 1 to 14 mW of laser power was used. Both power values were measured behind the objective lens. A 60X water-immersion objective lens (Olympus) was used for both photo-labeling and image acquisition. A GaAsP detector (Hamamatsu Photonics) was used for measuring green fluorescence. Photo-labeling was performed by drawing a region of interest — on average 1.0 x 1.0 µm — either in the center of the targeted glomerulus (for labeling projection neurons) or in the center of the soma (for labeling Kenyon cells). Photo-labeling projection neurons: Photoactivation was achieved through two to four cycles of exposure to 710- nm laser light, during which each pixel was scanned four times, with 25 repetitions per cycle, and 15 min rest period between each cycle. Image acquisition was performed with the laser tuned to 925 nm at a resolution of 512 by 512 pixels with a pixel size of 0.39 μm and a pixel dwell time of 4 μs; each pixel was scanned twice. A minimum of five samples per species were analyzed for each type of projection neuron. Photo-labeling Kenyon cells: Photoactivation was achieved through three to five single scans with the laser tuned to 710 nm, during which each pixel was scanned eight times. Before image acquisition, a 10 min rest period was implemented to allow diffusion of the photoactivated fluorophore within the neuron. Image acquisition was performed at a resolution of 512 by 512 pixels with a pixel size of 0.39 μm and a pixel dwell time of 4 μs; each pixel was scanned 2 times.

### Mapping Kenyon cell input connections using dye electroporation

The projection neurons connecting to a photo-labeled Kenyon cell were identified based on previously published protocols^25,28^. See Extended Data Figure 3 for a schematic depicting the procedure. In short, electrodes were made by pulling borosilicate glass pipette with filament (Sutter Instruments, BF100-50-10) to a resistance of 9-11 MΩ, fire-polished using a micro-forge (Narishige) to narrow their opening, and backfilled with 100mg/ml 3000-Da Texas-dextran dye (Thermo-Fisher, D3328). Under the guidance of a two-photon microscope (Bruker, Ultima), an electrode was centered into the post-synaptic terminal — or ‘claw’ — of a photo-labeled Kenyon cell using a motorized micromanipulator (Sutter Instruments). Short current pulses (each 10-50 V in amplitude and 0.5 millisecond long) were applied until the projection neuron connecting to the targeted Kenyon cell claw was visible. Not all the projection neurons connecting to a given Kenyon cells were dye-filled but on average 4 ± 1 of the claws formed by a given Kenyon cell were dye-filled. An image of the antennal lobe was acquired at the end of the procedure. Dye-labeled glomeruli were identified based on their shape, position and the location of their soma as defined in the available anatomical maps and the *Drosophila melanogaster* hemibrain connectome v1.2.1^13,14,29,41^.

### Dye-labeling individual projection neurons

Individual projection neurons were dye-labeled using a previously published protocol^16,43^. The cell body of the projection neuron of interest was first identified by lightly photo-labeling all the projection neurons innervating a given glomerulus by performing a single cycle of exposure to 710 nm light. After a rest period of 10 min, an unpolished electrode filled with Texas Red dextran dye was attached to the cell body of one of the photo-labeled projection neurons, and the dye was electroporated into the neuron using short current pulses; each pulse was 10 to 30 V in amplitude and 0.5 millisecond long. A resting period of about 30 min allowed the dye to diffuse throughout the neuron. Image acquisition was performed at a resolution of 512 by 512 pixels with a pixel size of 0.39 μm and a pixel dwell time of 4 μs; each pixel was scanned two times. A minimum of five samples per species were analyzed for each type of projection neuron.

### Quantifying morphological features of projection neurons

Representative images of projection neurons were projected at maximal intensity using the ImageJ/Fiji software^44^ (National Institutes of Health). Projection neurons were counted based on the number of photo-labeled cell bodies observed in the anterior or lateral or ventral clusters of the antennal lobe. Primary branches were defined as processes that emerge from the main axonal projection that traverses the calyx of the mushroom body. The length of the branches formed by a projection neuron and the number of forks were quantified using the ‘Simple Neurite Tracer’ plugin for ImageJ/Fiji software, while the surface area of the axonal arbors in the mushroom body calyx and lateral horn was calculated using the ‘ROI Manager’ and ‘Measure’ features of this software^45^. Total projection neuron bouton volume for a given sample was measured using Fluorender (University of Utah Scientific Computing and Imaging Institute; version 2.26.2^49,50^): boutons were traced using the ‘Paint Brush’ function. To distinguish boutons from the background, the ‘Edge Detect’ parameter was kept on and the ‘Edge STR’ was fixed at 0.505, while the selection threshold was adjusted to different values depending on signal intensity^46^. The ‘Physical Size’ value of the traced boutons was reported as total bouton volume.

### Similarity trees

Similarity trees, which were built using either the connectivity frequencies of glomeruli or the coding sequences of olfactory receptors, were generated using the ‘Traditional Search’ option in the Tree Analysis Using New Technologies (TNT) program (v.1.5)^47^. The cDNA sequences of *D. melanogaster* olfactory receptors were used as queries in BLASTN searches of *D. simulans* and *D. sechellia* cDNA sequences using the default settings of BLAST+v.2.13.0; *D. elegans* and *D. yakuba* cDNA sequences were used as outgroups^48^. Genomic regions were annotated using the Gnomon gene prediction method^49^. Coding sequence alignment of olfactory receptors was generated with MUSCLE, and the evolutionary history was determined with the ‘Maximum Likelihood’ method^50,51^. Sequences were edited manually to obtain the final alignments.

### Aversive learning paradigm

The aversive learning paradigm was designed based on a previously published protocol^52^. See Extended Data Figure 10 for a schematic depicting the procedure. Flies were collected a few hours before performing the protocol and housed in regular food vials before being tested. Groups of flies — containing between 60 and 100 individuals — were introduced in a T-maze training apparatus (CelExplorer Labs Co., TMK-501) that was attached to a flowmeter that kept a constant stream of 0.7L/min (Dwyer Instruments, 116011-01). During the training phase, flies were exposed to a first stimulus, henceforth referred to as the ‘conditioned stimulus +’ (CS+), at the same time as they were subjected to a regimen of electric shocks delivered by a stimulator (Grass Instruments Co, S48) for a period of one min (12 pulses of 90 V at a frequency of 0.2 Hz); shortly after, flies were allowed to rest for 45 s while exposed to ambient air before being exposed to a second stimulus, henceforth referred to as the ‘conditioned stimulus -’ (CS-), for one min without experiencing any electric shocks; flies were then allowed to rest for 45 s. The conditioned stimuli were either an odor dissolved in mineral oil or mineral oil alone (Sigma-Aldrich, M5904); the odors used were farnesol (1:1000 in mineral oil; Sigma-Aldrich, 43348), hexanoic acid (1:1000 in mineral oil; Sigma-Aldrich, 21529), 3-octanol (1:1000 in mineral oil; Sigma-Aldrich, 218405) or 4- methylcyclohexanol (1:1000 in mineral oil; Sigma-Aldrich, 153095). The training phase was performed either once (single training) or repeated six times with a 15-minute-long inter-training interval (spaced training). Between training and testing, there was a resting phase during which flies were housed in regular food vials and kept in the dark. During the testing phase, flies were given the choice to enter an arm of the T-maze perfumed with CS+ or the other arm perfumed with CS-. The performance index (PI) was calculated as follows: PI = (CS+ - CS-)/(CS+ + CS-). Each *n* reported in the data sets represents the average values obtained in a pair of reciprocal experiments; in reciprocal experiments, the stimuli used as CS+ and CS-were switched. Innate odor acuity was measured by allowing flies to choose between either a chamber perfumed with mineral oil or a chamber perfumed with an odor over the course of two minutes. Shock acuity was measured by allowing flies to choose between a chamber lined with a copper grid onto which 90V electric shocks were delivered every five seconds over the course of one minute. All experiments were performed at 23°C and 55%-65% relative humidity under dim red light.

### Statistical analyses

For the statistical analyses of the data shown in Table 1, Figure 3, 4, Extended Data Table 1, 3 and Extended Data Figure 3, 4 and 9, *p*-values were computed using the Mann-Whitney U test; statistical significance is indicated as *p* < 0.05 (*), *p* < 0.01 (**) and *p* < 0.001 (***). For the statistical analyses of the data shown in Figure 2 and Extended Data Table 2, *p*-values were computed using the Fisher’s exact test; to control for false positives, *p*-values were adjusted with a false discovery rate of 10% using a Benjamini-Hochberg procedure. For the statistical analyses of the data shown in Figure 5, *p*-values were computed using the sample t test.

